# The choroid plexus maintains ventricle volume and adult subventricular zone neuroblast pool, which facilitates post-stroke neurogenesis

**DOI:** 10.1101/2024.01.22.575277

**Authors:** Aleksandr Taranov, Alicia Bedolla, Eri Iwasawa, Farrah N. Brown, Sarah Baumgartner, Elizabeth M. Fugate, Joel Levoy, Steven A. Crone, June Goto, Yu Luo

## Abstract

The brain’s neuroreparative capacity after injuries such as ischemic stroke is contained in the brain’s neurogenic niches, primarily the subventricular zone (SVZ), which lies in close contact with the cerebrospinal fluid (CSF) produced by the choroid plexus (ChP). Despite the wide range of their proposed functions, the ChP/CSF remain among the most understudied compartments of the central nervous system (CNS). Here we report a mouse genetic tool (the ROSA26iDTR mouse line) for non-invasive, specific, and temporally controllable ablation of CSF-producing ChP epithelial cells to assess the roles of the ChP and CSF in brain homeostasis and injury. Using this model, we demonstrate that ChP ablation causes rapid and permanent CSF volume loss accompanied by disruption of ependymal cilia bundles. Surprisingly, ChP ablation did not result in overt neurological deficits at one-month post-ablation. However, we observed a pronounced decrease in the pool of SVZ neuroblasts following ChP ablation, which occurs due to their enhanced migration into the olfactory bulb. In the MCAo model of ischemic stroke, neuroblast migration into the lesion site was also reduced in the CSF-depleted mice. Thus, our study establishes an important and novel role of ChP/CSF in regulating the regenerative capacity of the adult brain under normal conditions and after ischemic stroke.

## Introduction

The cerebrospinal fluid (CSF) produced by the choroid plexus (ChP) has long been thought to be a crucial regulator of brain development^1^, homeostasis^2^, and disease^3^. In homeostasis, the CSF is thought to act as a hydromechanic shield^4^ and a regulator of electrolyte balance^5,6^ and waste clearance in the CNS^7^. The ChP, in addition to producing the CSF itself, also possesses a rich secretome containing hormone transporters, cytokines, neuroactive compounds and other signaling factors critical for brain development and physiological functioning^8–11^. In disease, both the ChP and CSF have been proposed to play a role in Alzheimer’s disease^12^, stroke^13^, TBI^14^, hydrocephalus^15^ and other pathologies. Previous studies suggest that the ChP/CSF may regulate neurogenesis in the adjacent subventricular zone both during developmental stages^16^ and in adulthood^17^, as well as under normal physiological conditions and injuries such as ischemic stroke. In the SVZ wall, proliferating neural stem cells (NSCs) produce adult-born neuroblasts (NBs), which continuously migrate into the olfactory bulb (OB)^18^. The CSF flow generated by the beating of ependymal cilia at the SVZ wall has been proposed to stimulate NSC proliferation^19^ and direct the migration of NBs within the SVZ ^20^ via ChP-secreted factors such as Slit2 through its chemorepellent property on neuroblasts ^21^. However, whether other factors in the CSF can either promote or inhibit the migration of newly born neuroblasts under physiological condition or pathological conditions is largely unknown.

Adult neurogenesis has been proposed as a promising treatment avenue for ischemic stroke. As the treatment window for ischemic stroke is currently limited (4.5 hours for rT-PA and 24 hours for thrombectomy)^22^, the majority of patients are not eligible for these procedures and may suffer poor functional outcomes. Cerebral ischemia is known to stimulate adult SVZ neurogenesis^23^ and divert adult-born neuroblast migration into the lesion site^24^. Work by our group^25–27^ and others^28^ demonstrates that enhancing these processes facilitates functional recovery at advanced time points after stroke, which is particularly promising due to the narrow acute treatment window for ischemic stroke. However, our ability to target adult neurogenesis therapeutically is limited by our lack of understanding of the CSF-borne factors that may regulate it, including those produced by the ChP.

Additionally, regulation of CSF volume is thought to be important in brain homeostasis, however, while the deleterious effects of overproduction and/or insufficient drainage of CSF are well-characterized in hydrocephalus^15^, little is known about the effects of reduced CSF production and the lack of CSF volume, which is often seen in human CSF leak patients^29^. Clinically, an important intervention targeting the ChP is surgical ChP removal, which is used in treating children with ChP tumors^30^, and ChP cauterization is under clinical trials for treating neonatal hydrocephalus in combination with endoscopic third ventriculostomy^31,32^. Despite their perceived critical role in both homeostasis and disease, both the ChP and CSF remain among the most understudied areas in the central nervous system (CNS)^3^. This gap in knowledge may be due to the lack of non-invasive tools allowing the manipulation of normal CSF production and ChP function *in vivo*.

First published in 2005, the ROSA26iDTR mouse line^33^, a useful tool for cell ablation, saw widespread use in over 341^34^ studies to date to dissect out roles of specific cell types in both CNS and other organs^35^. Surprisingly, we discovered a previously unreported robust phenotype of loss of ventricular spaces upon diphtheria toxin (Dtx) administration that occurs in the ROSA26iDTR line in a Cre-independent manner^36^. This phenotype is of particular importance as it not only may have significant implications on the interpretation of previous studies utilizing this ROSA26iDTR mouse line^35^, but also provides a novel method to manipulate CSF volume in the adult mouse brain. However, the underlying mechanism behind this loss of ventricular spaces was previously not clear. In the current study, we demonstrate that the floxed R26-loxP-STOP-loxP-DTR cassette is “leaky” specifically in the choroid plexus epithelial cells, which accounts for apoptosis of these cells and subsequent loss of ventricular spaces in the iDTR mice after Dtx treatment. More importantly, using this tool, we directly investigate the role of ChP function in maintaining ventricular CSF volume and general brain health. Specifically, we characterize the first non-invasive and inducible mouse genetic model to assess the consequences of loss of ChP/CSF and its related-factors on survival, gross motor behavior and cognitive function. Furthermore, we directly examine the role of the ChP/CSF in maintaining the SVZ ependymal cilia structure and the neurogenic potential of SVZ under normal conditions and after ischemic stroke in the adult mouse brain.

## Results

### 1. CSF-producing choroid plexus epithelial cells express diphtheria toxin receptor (DTR) independent of Cre expression, and undergo apoptosis in response to Dtx in ROSA26iDTR mice

The ROSA26iDTR line (referred to as iDTR mice in this study, Jax #007900) was originally developed^33^ to enable cell-type-specific ablation in mice. Specifically, it carries a floxed transcription stop-cassette preceding the gene for the simian Heparin-binding EGF-like growth factor, HB-EGF, which acts as a receptor (DTR) for *C. diphtheriae* toxin (Dtx). Cre-driven excision of the stop cassette induces cell-type specific DTR expression sensitizing cells to the toxin. Upon binding of Dtx to DTR and its subsequent internalization, apoptosis is induced through the arrest of protein synthesis, resulting in cell type-specific ablation^38^. We initially reported unexpected and pronounced loss of ventricular volume in the ROSA26iDTR mouse line upon administration of Dtx but the underlying mechanism was not clear ^36^.

Considering that the phenotype described above could present a unique opportunity to examine the role of the ChP and CSF *in vivo* in the adult brain, we investigated the underlying mechanism behind CSF loss in this model.

Since the ChP has long been considered to be the main CSF-producing organ^6^, we hypothesize that the loss of CSF could be due to the Dtx-induced apoptosis in the CSF-producing ChP epithelium which might have “leaky” expression of the DTR gene in a Cre-independent manner. To test whether DTR could be expressed in the ChP or other brain regions and peripheral tissues in iDTR mice, we harvested RNA from ChPs of three cerebral ventricles and a variety of brain regions as well as some peripheral epithelial organs (skin from the ear, lung, and small intestine) from WT, iDTR heterozygous, and homozygous mice (none of which express Cre recombinase). Indeed, using quantitative reverse transcription-PCR (qRT-PCR), we observed that a substantial amount of DTR mRNA is transcribed in iDTR heterozygous and homozygous mice (Fig. 1 B), particularly enriched in the ChP from the lateral ventricle (LV), 3^rd^ ventricle (3V) and 4^th^ ventricle (4V). The quantity of DTR RNA in the ChP generally shows higher levels in homozygous iDTR mice than heterozygous mice with a similar expression pattern among tested regions/organs. Interestingly, the levels of DTR expression in ChP are much higher than in various brain parenchymal regions (Fig 1B) and peripheral epithelial tissues (Fig 1 B), with the gut having the highest level amongst the peripheral tissues examined, but still substantially lower than ChP levels. Importantly, all RNA samples were treated with DNase, and RT(-) control shows no amplification in the qPCR reaction, validating that the RNA levels of DTR detected in our study are not due to DNA contamination. As expected, DTR expression was not detectable in any of the WT tissues examined. To further validate the presence of DTR in the ChP of iDTR mice, we performed immunohistochemistry to detect DTR protein with an antibody specific for simian and not mouse HB-EGF and only detected DTR immunoreactive signal in the ChP but not from other brain regions in the iDTR mice (Fig. 1 C, D). Next, we examined whether apoptosis occurs in the ChP of iDTR mice after Dtx administration. To this end, we performed TUNEL staining on day 3 after the first Dtx injection (20ng/g/day), which showed extensive apoptosis within the ChP in iDTR + Dtx cohort but not in WT + Dtx or iDTR + saline cohort (Fig. 1 E-G). In WT + Dtx or iDTR + saline controls, sparse TUNEL positive cells were only detected in the regions where ongoing adult neurogenesis is present (the subventricular zone – SVZ and the subgranular zone – SGZ) consistent with previous reports^39,40^. However, in the iDTR + Dtx mice, in addition to the sparse TUNEL+ cells in the SVZ and SGZ regions, substantial TUNEL positive cells were also specifically detected in the ChPs. To examine which cell types in the ChP undergo apoptosis after Dtx treatment (Fig. 1 H – J), we performed co-immunostaining for the α1 subunit of Na+/K+ ATPase (marker for the apical surface of the ChP epithelial cells), CD68 (macrophage marker), IB4 (blood vessel endothelium) and PDGFRβ (blood vessel pericytes). We determined that compared to other cell types in the ChP, apoptotic cell death occurred predominantly (93.5±5.7%) in α1-positive ChP epithelial cells (Fig. 1 K), indicating that it is the loss of CSF-producing ChP epithelial cells that leads to the loss of CSF volume and ventricular spaces in iDTR mice after Dtx treatment.

**Figure 1.**
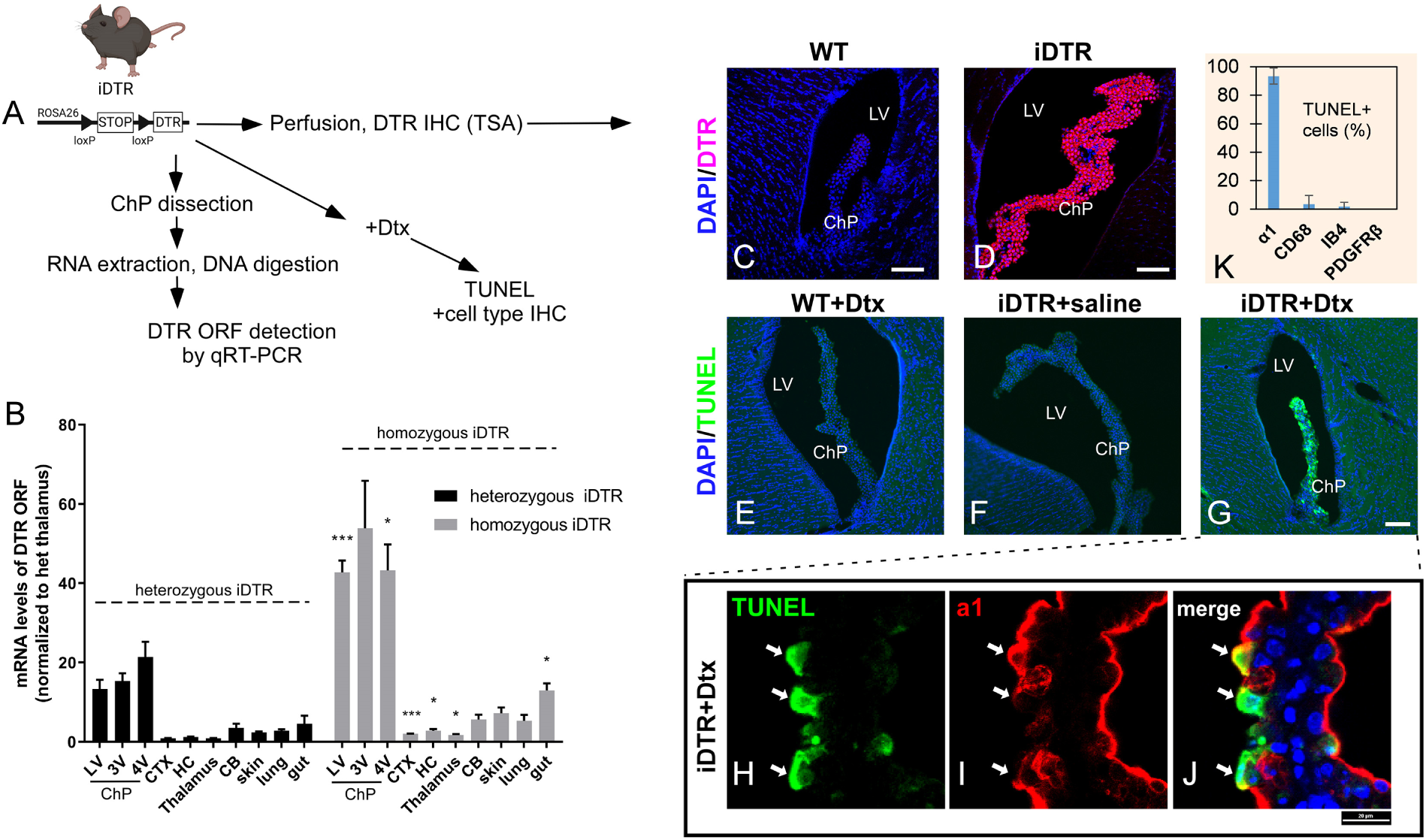
Choroid plexus (ChP) epithelial cells undergo apoptosis in response to Dtx treatment in iDTR mice. (A) Schematic of experimental paradigm. (B) Expression levels of DTR ORF mRNA in the ChP and brain and peripheral tissues of 5-7 month-old iDTR^+/WT^ (heterozygous) and iDTR^+/+^ (homozygous) mice (mean ± SEM, n= 3-4 per tissue per genotype, Student’s t-test or nonparametric equivalent between genotypes within each tissue). CTX – cortex, HC – hippocampus, CB – cerebellum. No DTR expression was detected in WT mice (data not shown). (C, D) –presence of DTR protein (red) in the ChP of iDTR (D) but not WT mice (C). (E-G) TUNEL staining (green) showing apoptosis specifically in the ChP of Dtx-treated iDTR mice (G), but not Dtx-treated WT mice (E) or saline-treated iDTR mice (F), scale bar - 100 µm. (H-I) Magnified confocal image of a lateral ventricle ChP of a Dtx-treated iDTR animal stained with TUNEL (green, H), and ChP epithelium cell marker (α1 subunit of Na^+^/K^+^ ATPase, red, I) and merged image (J). (K) – quantification of TUNEL+ cells that co-express different cell type markers (α1 – apical ChP epithelial cells, CD68 – macrophages, IB4 – endothelial cells, PDGFRβ – pericytes) in the ChP of Dtx-treated iDTR animals (mean ± SD, n = 3. Scale bar – 20 µm). * p < 0.05, ** p < 0.01, *** p < 0.001.

Next, we further quantitatively evaluated the degree of ChP epithelial cell ablation following different dosages of Dtx administered to iDTR mice. Adult iDTR heterozygous mice were subjected to three different dosages of Dtx (20ng/g/day for 3 days, 4ng/g/day for 3 days, 10ng/g/day for 2 days) and a control group of WT mice were treated with 20ng/g/day for 3 days. Brain tissue was collected at one month post-Dtx injection (a time point where we observed that the loss of ventricular volume is stable after the initial Dtx injection) to evaluate extent of ChP epithelial cell loss. First, we collected serial coronal sections and quantified ChP area by transthyretin (TTR) immunostaining, which is an additional marker for the entire ChP epithelium (Fig. 2 A-I). We observed a marked decrease in TTR+ ChP epithelium area with all dosages of Dtx to a similar extent in the 3^rd^ ventricle ChP, while in the lateral ventricle ChP the 20 ngX3 dosage produced the greatest degree of ablation (Fig. 2 J). However, considering that the plane of coronal sections could obscure changes occurring along the entire length of the ChP, we extracted entire lateral ventricle ChPs from a cohort of WT and iDTR mice one month post-ablation and performed ChP whole mount immunohistochemistry to stain for TTR (ChP epithelial cells) and CD31 (blood vessel endothelium marker). Consistent with the coronal section evaluation, ChP whole mount immunostaining also shows (Fig. 2 K-M) that the total surface area of ChP is decreased following the high (20ng/g/day X3) more so than the low (4ng/g/dayX3) dose of Dtx and, interestingly, loss of ChP epithelial cells and shrinkage of total ChP surface area resulted in to increased blood vessel linear length normalized to ROI area (defined as blood vessel length density in our study) in ChP-ablated mice (Fig. 2 N). This suggests that loss of ChP epithelial cells might lead to secondary changes in other cell types in the ChP, which warrants further investigation in future studies. At 3 or 6 months post-Dtx treatment, we observe almost complete absence of ChP in iDTR mice, therefor ChP whole mounts could not be collected at these time points to evaluate the remaining ChP size. Interestingly, this finding demonstrates the absence of regeneration of the adult ChP epithelium after ablation, which has not been specifically tested previously.

**Figure 2.**
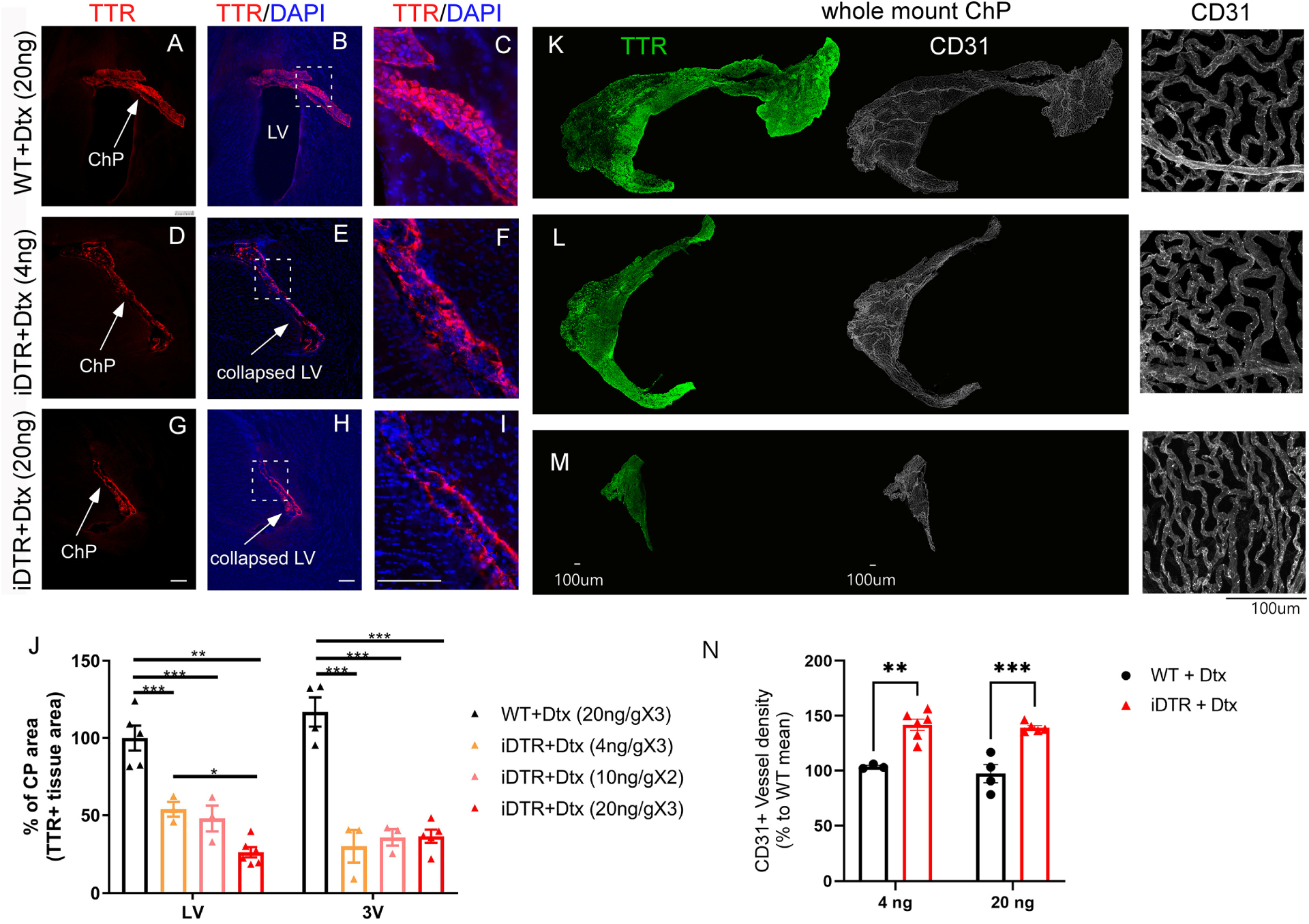
Dose-dependent loss of ChP area in response to Dtx treatment in iDTR mice. (A-I) Representative images of coronal sections showing ventricle and the ChP stained for TTR (red) and DAPI (blue) 2-5 weeks after Dtx treatment, 3-5 month-old animals. LV – lateral ventricle. Scale bar – 100 µm. (C, F, I) Magnified portions of B, E, H to show loss of TTR+ cells and the ventricle space in Dtx-treated iDTR mice. (J) Quantification of TTR+ area in coronal sections, normalized to WT mean (n = 3-6 per group, mean ± SEM, One-way ANOVA). (K-M) Whole mount ChPs at one month post-Dtx stained for TTR (ChP epithelial cells, green) and CD31 (blood vessel endothelium, white), 2-month-old animals. Right side – magnified portion of the whole mount to show blood vessel density. (N) Quantification of blood vessel length density (normalized to WT mean, n= 3-6 per group, mean ± SEM, Two-Way ANOVA). * - p < 0.05, ** - p < 0.01, *** - p < 0.001.

Thus, we demonstrate that administration of Dtx to iDTR mice is a non-invasive way to selectively ablate the CSF-producing epithelial cells. Taking advantage of this model, we investigated the role of ChP/CSF in general brain health and some key neurological processes that have been long speculated to be dependent on normal ChP/CSF functions.

### 2. Rapid loss of ventricular spaces in Dtx-treated iDTR mice following ChP ablation

To examine the timeline of CSF loss following ChP ablation, we administered 20 ng/g/day of Dtx over three days in young adult mice (3 - 4 months old, female and male animals, Supplementary Fig. 1 A). The loss of ventricle volume was visible on T2-weighted MRI scans as early as three days after the first Dtx injection (Supplementary Fig. 1 B). Both female and male iDTR mice showed a similar pattern and extent of CSF loss following ChP ablation. The reconstruction of ventricular volume from T2-weighted slices showed a pronounced decrease in the volume of lateral and third ventricles on day 3 after start of Dtx treatment in iDTR but not in WT mice. (Supplementary Fig. 1 C). The ventricular size phenotype was also observed in PFA-fixed histologically processed coronal brain sections (Supplementary Fig. 1 D - G). These results confirm our previous report^36^ that the loss of ventricular space occurred in a Cre-independent manner, demonstrating that only the ROSA26iDTR allele is required to induce ventricle loss. Next, to further investigate how fast CSF is depleted in Dtx-treated iDTR mice, we administered Dtx at 20 ng/g/day X 3 to 3–4-month-old mice and acquired T2-weighted MRI scans one day before the start of Dtx treatment and at one, two, and three days after the first Dtx injection (Fig. 3 A). In Dtx-treated iDTR mice, the loss of ventricular volume has already initiated by day 1 after the start of Dtx treatment, exacerbating by day 2 and proceeding to almost complete loss of ventricular volume on day 3 post-Dtx (Fig. 3 B, C). That suggests either the requirement of at least three consecutive Dtx doses or 3 days for the apoptosis in the ChP epithelium to reach its peak or it could also reflect the delayed relationship between the loss of ChP epithelium and the loss of CSF, albeit the latter appears less likely due to rapid rates of CSF circulation in the brain. Furthermore, to validate the loss of CSF volume seen on MRI, we collected CSF from the cisterna magna (CM) of WT and iDTR mice post-Dtx (Fig. 3 D – F). On average, WT mice yielded 7.3 µl of CSF, while the iDTR mice only yielded 1.66 µl. Additionally, the CSF volume loss was visually apparent due to the flat, collapsed appearance of the CM in iDTR mice, as opposed to WT mice (Fig. 3 D, E). Given the potentially broad impact of this phenotype on numerous prior studies^34^, to validate this as an authentic phenotype, an independent lab (Goto lab) replicated these findings from iDTR breeders recently purchased from Jackson Laboratories. Additionally, we validated these findings in tissue sections from our previous experiments utilizing the ROSA26iDTR line with or without Cx3cr1^CreER^ (expressed primarily in peripheral macrophages and microglia), Glast^CreER^ (expressed in astrocytes) or mGFAP-Cre lines (expressed primarily in astrocytes and neural stem cells) maintained and treated with Dtx in the Luo lab (Supplementary Fig. 2). Moreover, retrospectively examining tissue preserved in 2015-2017 from a third independent lab (Crone lab), we observed the same phenotype of loss of ventricular spaces in the Dtx-treated ROSA26iDTR mice with or without Chx10^Cre^ (in this line^41^, Cre expression is restricted to the spinal cord, brainstem, and the eye but absent in the forebrain) (Supplementary Fig. 2). Thus, multiple independently maintained iDTR mouse lines dating back to at least 2015 demonstrate loss of ventricular volume following Dtx treatment, independent of Cre expression.

**Figure 3.**
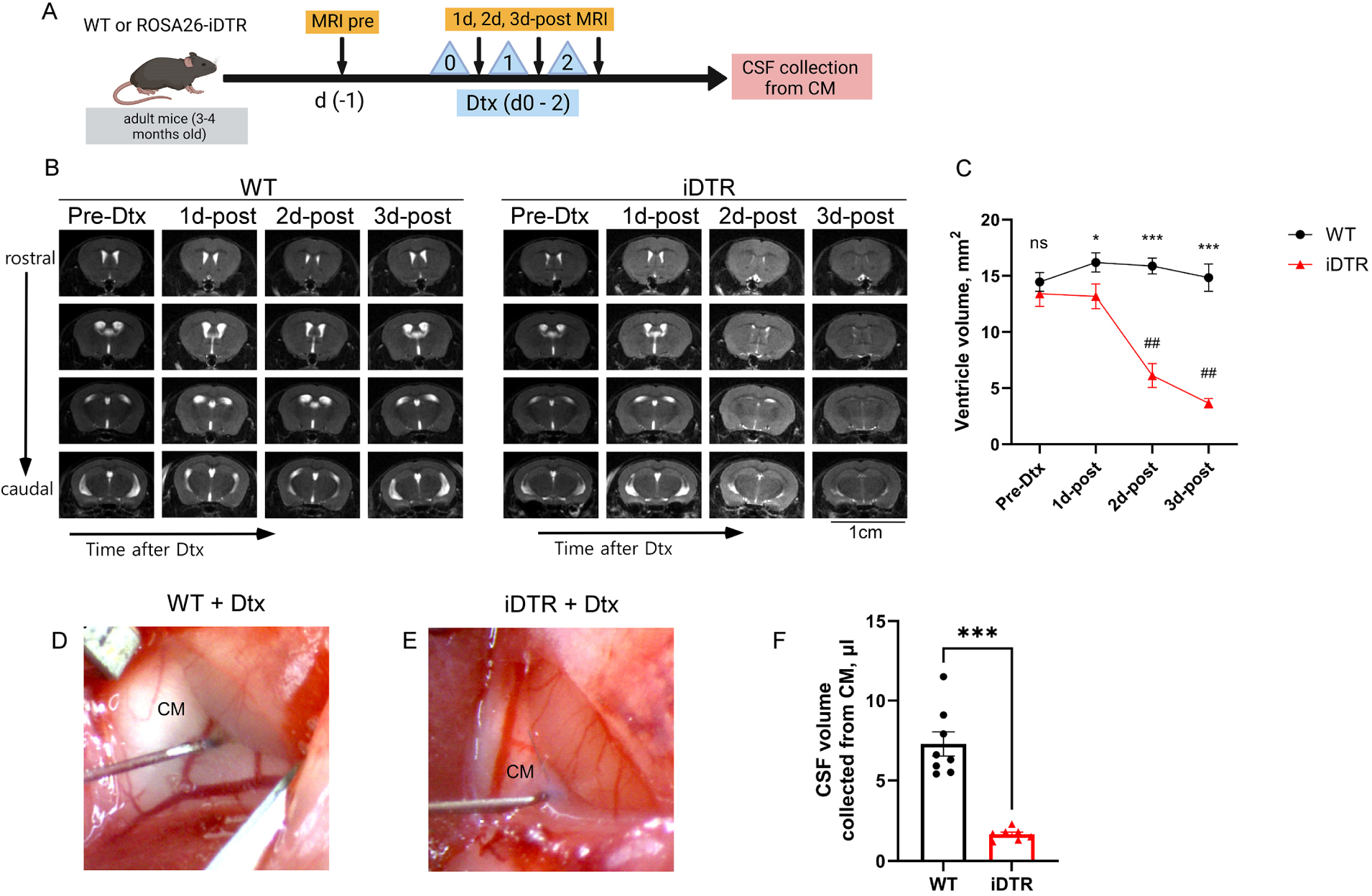
Time course of CSF volume loss after Dtx treatment. (A) Schematic of experimental design to track daily CSF volume change. (B) T2-weighted MRI showing ventricular spaces in WT (left side) and iDTR (right side) brain. Scale bar – 1 cm. (C) – planimetric quantification of ventricle volume from 9 T2 MRI slices in WT and iDTR mice one day before Dtx, and at 1, 2, and 3 days post-Dtx, n = 4 per group, mean ± SEM, Two-Way ANOVA with genotype and timepoint as factors. Comparisons between genotypes: ns – p ≥ 0.05, * - p < 0.05, *** - p < 0.001. Comparisons within genotype to pre-Dtx volume: ## - p < 0.01. (H, I) – example images of the cisterna magna (CM) of WT (D) and iDTR (E) mice at 6 months post-Dtx during CSF collection. (F) – quantification of the volume of CSF collected from the CM (mean ± SEM, n = 7-8 per group, Mann-Whitney U test). *** - p < 0.001.

#### Lateral and 3^rd^ ventricles experience more profound volume loss compared to 4^th^ ventricle and pineal recess after Dtx treatment

To quantify the CSF volume loss in each ventricle *in vivo*, we developed a T2 fluid-sensitive MRI sequence that allows for the detection of CSF in live animals and reconstruction of 3D CSF-filled ventricular spaces in mouse brain. Utilizing this sequence, we examined the distribution of CSF volume in cohorts of WT and iDTR mice treated with either 4 or 20 ng/g/day of Dtx for three consecutive days at 5-18 days after Dtx treatment. As seen in 3D reconstructions of the T2 fluid-sensitive MRI data (Fig. 4 A), the lateral and third ventricles undergo profound volume loss upon Dtx administration in iDTR mice, with 4 ng/g/day exerting a partial effect compared to 20 ng/g/day (Fig 4 B). The 4^th^ ventricle and the pineal recess seem to be less susceptible to CSF volume loss, with 4 and 20 ng/g/day affecting their volume to a similar extent (Fig 4 B).

**Figure 4.**
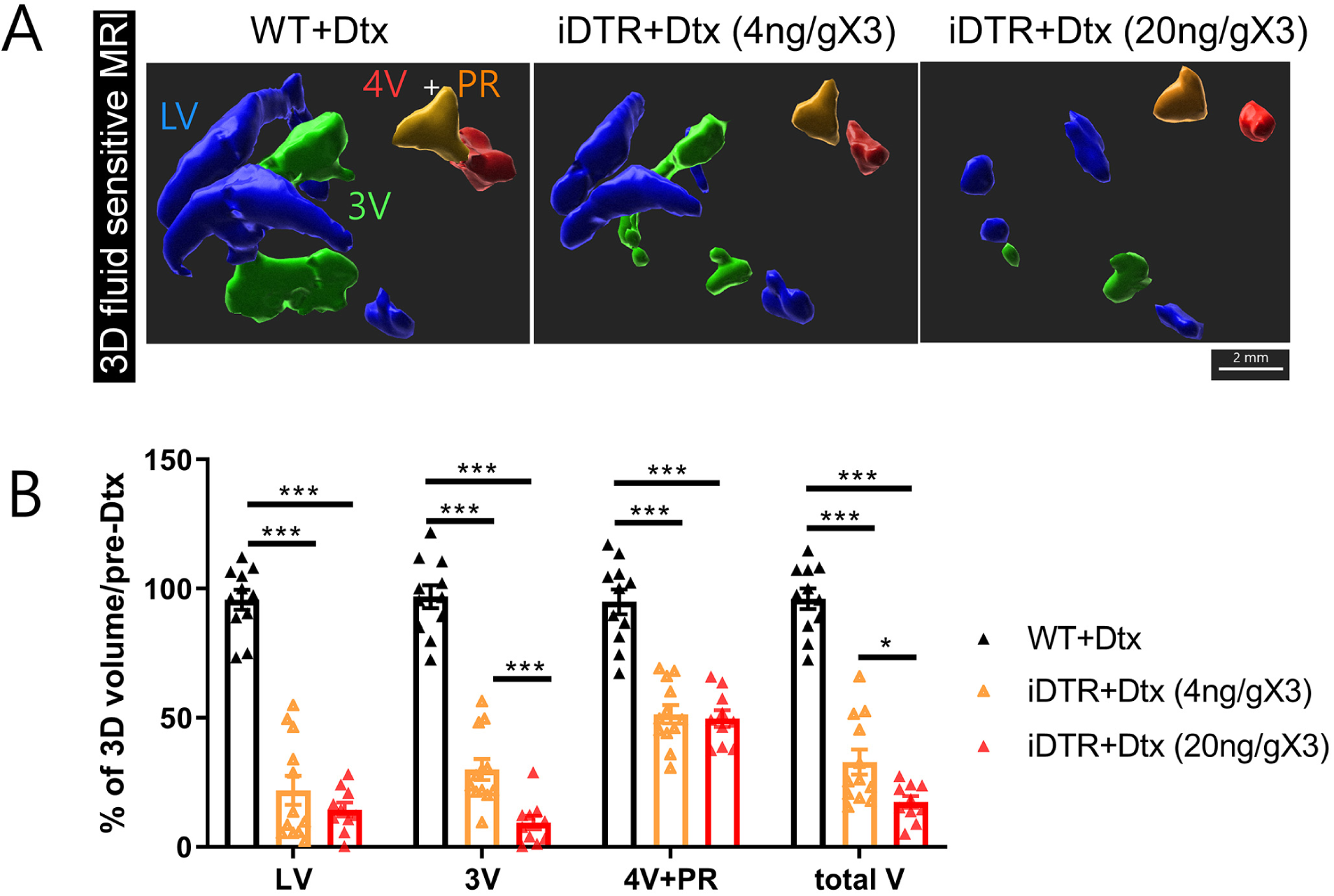
Dose-dependent CSF volume loss after Dtx treatment. Top panel – representative 3D reconstructions of ventricular spaces from T2 fluid-sensitive MRI data acquired one day before Dtx treatment and 5-18 days after the start of Dtx treatment in 2-10 month-old animals. LV – lateral ventricles (blue), 3V – third ventricle (green), PR – pineal recess (orange), 4V – fourth ventricle (red). Scale bar - 100 µm. Bottom panel – quantification of ventricular volume from 3D reconstructions of T2 fluid-sensitive MRI data (normalized to pre-Dtx mean ventricular volume, n = 10-12 per group, mean ± SEM, One-Way ANOVA) * - p < 0.05, ** - p < 0.01, *** - p < 0.001, **** - p < 0.0001.

#### Stable and sustained loss of CSF volume after ChP ablation

Next, to investigate whether the loss of CSF volume was sustained in this model, we administered 20 ng/g/day of Dtx for 3 days to cohorts of 2-month-old WT and iDTR mice and aged them for up to 20 months post Dtx. Using the T2 fluid-sensitive MRI sequence, the animals were scanned before Dtx administration, at day 3 after the start of Dtx treatment and at 8 and 20 months post-Dtx (Supplementary Fig. 3 A, B). The rapid and profound loss of ventricular volume observed at day 3 post-Dtx was present for up to 20 months (the longest timepoint we tested). The lateral and third ventricles exhibited a nearly total loss of CSF (Supplementary Fig. 3 C, D), while the 4^th^ ventricle and the pineal recess displayed a smaller decrease in CSF volume (Supplementary Fig. 3 E). These results indicate that alternative sources of CSF production or regeneration of ChP after ablation do not occur or are not sufficient to restore ventricular volume. Over the span of 20 months in this small cohort (n=3-4 per group), we did not observe any unusually high mortality or gross motor abnormalities in iDTR mice that received Dtx.

#### CSF volume reduction after ChP ablation is observed in both young (1 month) and old (20-24 month) iDTR mice treated with Dtx

Evidence from human and rodent studies shows profound changes in the production rate^42^ and composition^43,44^ of the CSF with aging^45^, which has been linked to numerous pathological processes^1,15,46,47^. Considering the potential relevance of this model to the role of CSF in aging we sought to determine whether the capacity of Dtx to induce CSF volume loss in iDTR mice changes with age. To this end, we administered Dtx (20 ng/g/day x 3) to cohorts of young (1 month old) and old (20-24 month old) WT and iDTR mice. On T2 fluid-sensitive MRI scans performed on day 3 after the first Dtx injection, we observed a similarly pronounced loss of CSF volume in both cohorts, as well as a similar pattern of CSF volume loss by ventricle in young mice (Supplementary Fig. 4 A) and aged mice (Supplementary Fig. 4 B).

### 3. ChP/CSF loss does not cause an acute neurobehavioral phenotype

Unexpectedly, ChP ablation and subsequent loss of CSF after Dtx treatment did not lead to gross and acute neurological and motor deficits in our model. In fact, ChP-ablated animals seem to survive for up to 20 months post-Dtx (Supplementary Fig. 3) without any visually apparent differences from age-matched WT controls, including in body weight at day 4, 7, and 30 post-Dtx (short-term), as well as at 5 months post-Dtx (long-term) in a separate cohort of animals (Fig. 5 A). To investigate whether ChP ablation could induce a more subtle neurological phenotype, we examined the general behavioral functioning of the animals at 1 month post ablation (Fig. 5 B – H). We conducted a comprehensive battery of tests to examine general motor function, circadian locomotor activity rhythms, and cognitive function in two different cohorts of ChP-ablated mice given either 4 ng/g/day x 3 (Fig 5 B, D, F) or 20 ng/g/day x 3 (Fig. 5 C, E, G, H) of Dtx. At one-month post-ablation, ChP-ablated mice exhibited similar locomotor activity in a 23-hour time period (Fig. 5 B, C), as well as a normal light-dark circadian locomotor activity rhythm (Fig. 5 D, E). Cognitive function measured either by Barnes Maze (Fig. 5 F) or spontaneous alternation Y-maze (Fig. 5 G, H) showed no differences at 1 month post ChP ablation. These findings in themselves are surprising given the numerous essential homeostatic functions ascribed to the CSF and the ChP^6^, such as maintenance of electrolyte levels^5^ that are essential for sustaining and generating membrane potentials, as well as carrying essential hormones^10^ and neuromodulators^11^ and possibly removing waste products^48,49^, as well as due to the fact that some of the key brain regions responsible for motor control (striatum and cerebellum), or cognitive function (hippocampus), lie in close proximity to the ventricular system.

**Figure 5.**
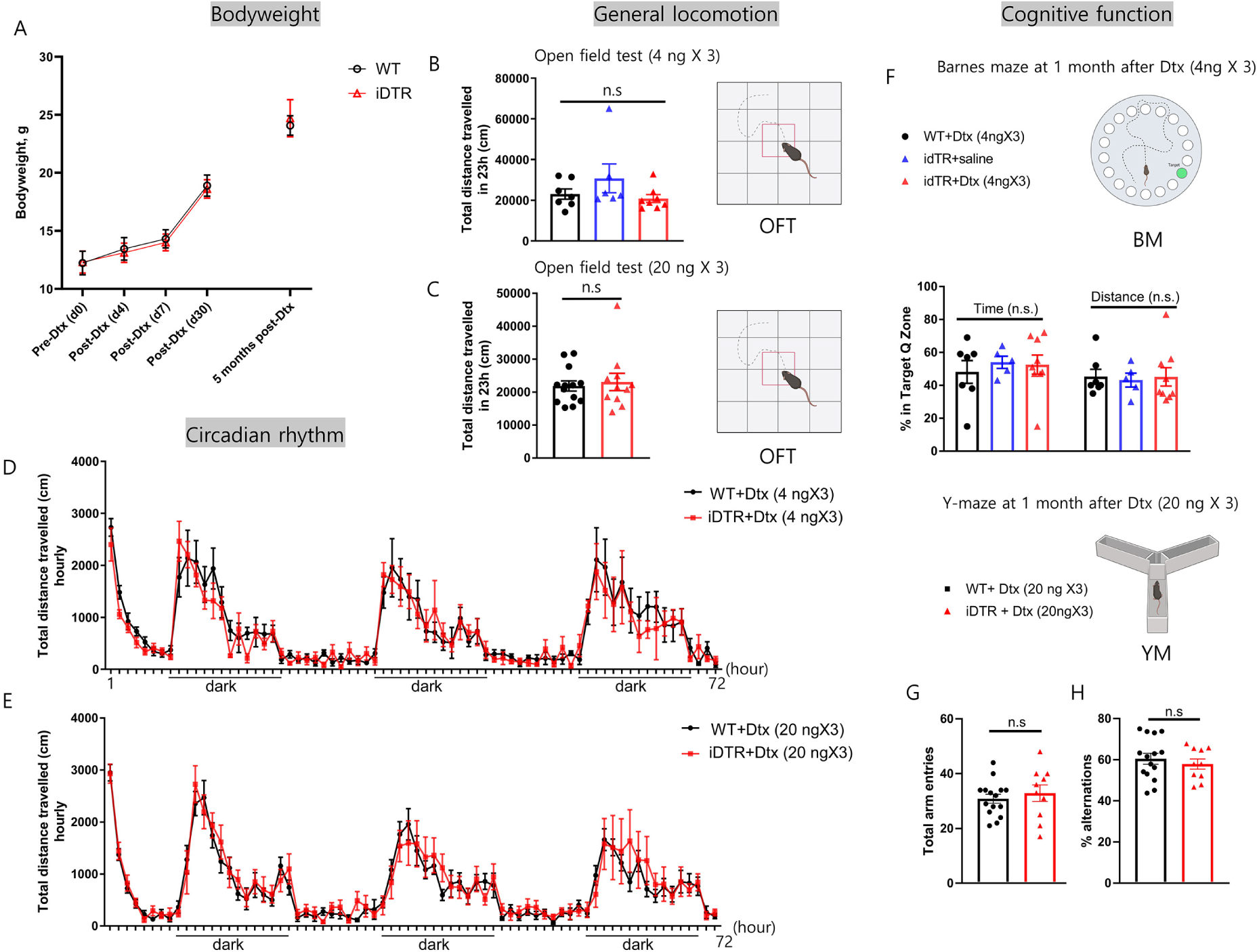
Behavioral characterization of ChP-ablated mice at 1 month post Dtx treatment. (A) – change in body weight in a cohort of animals treated with Dtx at P30, followed up to day 30 post-Dtx, and a separate cohort weighed at 5 months post-Dtx. (B, D, F) – behavioral data from a cohort treated with 4 ng/g/day x 3, n = 6-8 per group. (C, E, G, H) – data from a cohort treated with 20 ng/g/day x 3, n = 10-15 per group. (B, C) – total distance travelled in a 23hr period in an automated open field paradigm. (F) – percentage of time spent, and distance travelled in the target quadrant in Barnes maze. (G, H) – total arm entries (G) and alternation rate (H) in a spontaneous alternation Y-maze. (E, F) – hourly motor activity over a 72 hr period in the automated open field test. Experiments conducted in 2-4 month-old animals. Data presented as mean ± SEM. (A) – Tw-Way RM ANOVA with timepoint and genotype as factors. B, F – One-way ANOVA. C, G, H – Student’s t-test or Mann-Whitney U test. Data are presented as mean ± SEM. n.s – p ≥ 0.05.

### 4. The choroid plexus/CSF are essential for newly-born neuroblast retention in the SVZ

#### Choroid plexus ablation does not affect the Ki67+ proliferating cell number but leads to reduced newly born neuroblast (doublecortin, DCX+) number in the SVZ in the adult mouse brain

Given the close proximity of the subventricular zone (SVZ) neural stem cells to the CSF and the lateral ventricle ependymal cell layers, we took advantage of this model to evaluate the long-proposed role of CSF in regulating adult neurogenesis^17,20,21,50,51^. WT and iDTR heterozygous mice were treated with 4 or 20 ng/g/dayX3 dosage regimen of Dtx. Doublecortin (DCX) was used as a marker for newly born neuroblasts and Ki67 as a marker for proliferating cells. For DCX and Ki67 quantification, the animals were treated with Dtx and harvested at 4 weeks (Supplementary Fig. 5) or either 3 months or 6 months post Dtx injection (Fig. 6). Previous studies have shown that the SVZ newly born neuroblasts can be added into the olfactory bulb between 2-4 weeks after their birth as new granule neurons^18,52^, we therefore first evaluated an earlier timepoint of 4 weeks post-ablation (Supplementary Fig. 5) by immunostaining for the newly born neuroblast cell marker (DCX) in coronal sections containing the SVZ. The high (20 ng/g/day, Supplementary Fig. 5 B, b’) dose of Dtx resulted in a significant decrease in DCX+ area fraction in iDTR mice compared to WT mice (Supplementary Fig. 5 A, a’), albeit characterized by considerable variability (Supplementary Fig. 5 C). To test whether this initial trend of decrease in the SVZ neuroblast pool will be sustained over time, we evaluated a cohort of animals at 3 months post Dtx by immunostaining for neuroblasts (DCX) and proliferating cells (Ki67) (Fig. 6 A-F). Previous studies have suggested that ChP provides factors in the CSF that are critical for maintaining the proliferation of SVZ neural stem cells and rapidly dividing progenitor cells^16,51^, however, our results show that at 3 months post ChP/CSF ablation, the total Ki67+ cell number was not affected by ChP ablation in iDTR mice (Fig. 6 C, E). DCX immunoreactive area in the dorsal corpus callosum, which is not in direct contact with the CSF was not affected in the 3-month post-Dtx group (Fig. 6 D). However, in the ventral SVZ which directly contacts the CSF, DCX immunoreactive area was significantly decreased in ChP-ablated mice, indicating a reduced number of DCX+ newly born neuroblasts (Fig. 6 F). Since ChP ablation leads to shrinkage of lateral ventricles which might affect the overall shape or the anterior-posterior position of the coronal section, we further examined the SVZ in whole mount preparations in both 3-month and 6-month ChP-ablated mice compared to Dtx-treated WT mice (Fig. 6 G-L). Strikingly, the results from SVZ whole mount immunostaining further validate the absence of an effect of ChP ablation on Ki67+ cell density in the lateral wall of the SVZ (Fig. 6 I, L). However, consistent with the results from coronal section quantification, SVZ whole mounts show significantly decreased DCX immunoreactive area in iDTR mice both 3 and 6 months after Dtx treatment (Fig. 6 H, K). This decrease in DCX+ cells in the SVZ whole mounts was not observed in age-matched iDTR mice that did not receive Dtx treatment (data not shown). Additionally, as expected, we observed a dense migratory network of DCX+ cells in WT SVZ whole mounts, which is mostly absent in the SVZ of ChP-ablated mice. Our results indicate that ChP/CSF-derived factors might not be critical for maintaining the proliferative potential of SVZ neural stem cells and/or transient amplifying progenitors but instead are critical for the maintenance of neuroblast number in the SVZ and/or the migration within the network formed by DCX+ neuroblasts.

**Figure 6.**
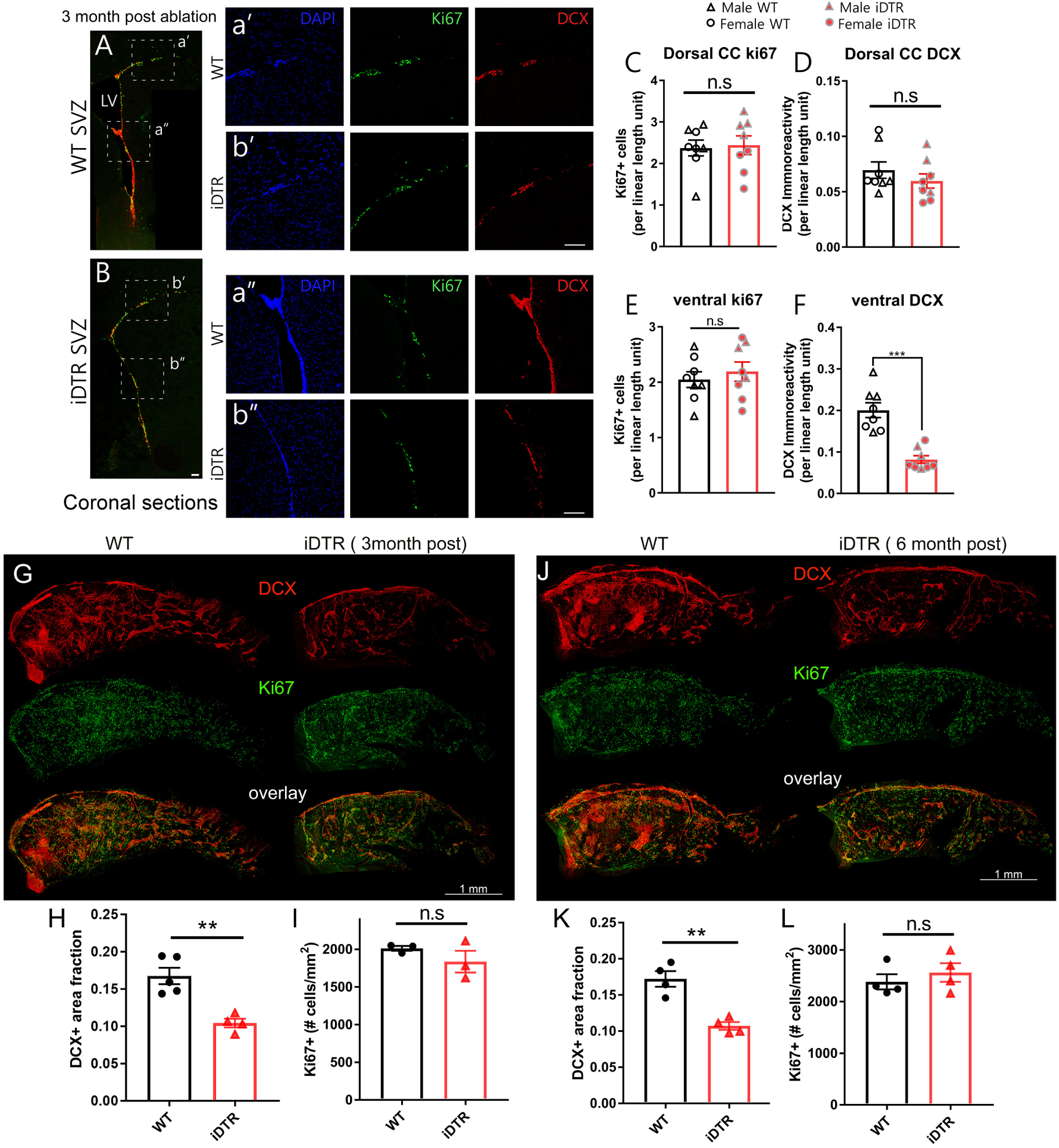
Proliferating cells and newly-born neuroblasts in the SVZ at 3 and 6 months post-Dtx. (A-F) – data from coronal tissue sections at 3 months post Dtx. (A, B) – representative scans of coronal sections containing the SVZ area and the dorsal corpus callosum. Scale bar – 100 µm. LV – lateral ventricle. (a’, a’’, b, b’’) – magnified portions of A and B, containing the dorsal corpus callosum (a’, b’) and the SVZ (a’’, b’’) stained for DAPI (blue), Ki67 (green), DCX (red). (C, E) – quantification of ki67+ cells (normalized to length) in the dorsal corpus callosum (C) and the ventricular wall. (D, F) – quantification of DCX+ area (normalized to SVZ length) in the dorsal corpus callosum (D) and the ventricular SVZ (F). Data are mean ± SEM, n = 8 per group, Two-way ANOVA with sex and genotype as factors. No statistically significant sex differences were detected. (G-L) – data from SVZ whole mount preparations at 3 months (G-I) and 6 months (J-L) post Dtx. Scale bar – 1 mm. (G, J) – representative SVZ whole mounts for the 3 month cohort (G) and the 6 month cohort (J) stained for DCX (red) and ki67 (green). (H, K) – DCX+ area fraction at 3 and 6 months post ablation. (I, L) – Ki67+ cell density at 3 and 6 months post ablation. Animals were 6-7 months old at the time of tissue collection. Data are mean ± SEM, n = 3-4 per group, Student’s t-test or Mann-Whiney U test). n.s – p ≥ 0.05, ** - p < 0.01, *** - p < 0.001.

Previous reports suggest that the directional flow of CSF and the polarity of motile cilia of the ependymal cells lining the lateral ventricular wall might be critical for neuroblast migration in the SVZ^21^ (Fig. 7 A). We therefore further hypothesize that the loss of CSF-borne factors and overall CSF circulation could affect the motile cilia of SVZ ependymal cells as well, which could lead to disruption of the migratory network and reduced neuroblast number in the SVZ after ChP ablation. We stained SVZ whole mount preparations at 3 months post ablation for acetylated tubulin (Fig. 7 B-E) which visualizes motile cilia and their polarity and direction. Interestingly, our results show that in ChP-ablated mice, we observed a disorganized pattern and loss of the directional orientation of acetylated tubulin-positive bundles of motile cilia, which could underlie the differences in DCX+ neuroblast number in the SVZ, as well as the altered migratory network that we observed in ChP-ablated mice (Fig. 6 G, J).

**Figure 7.**
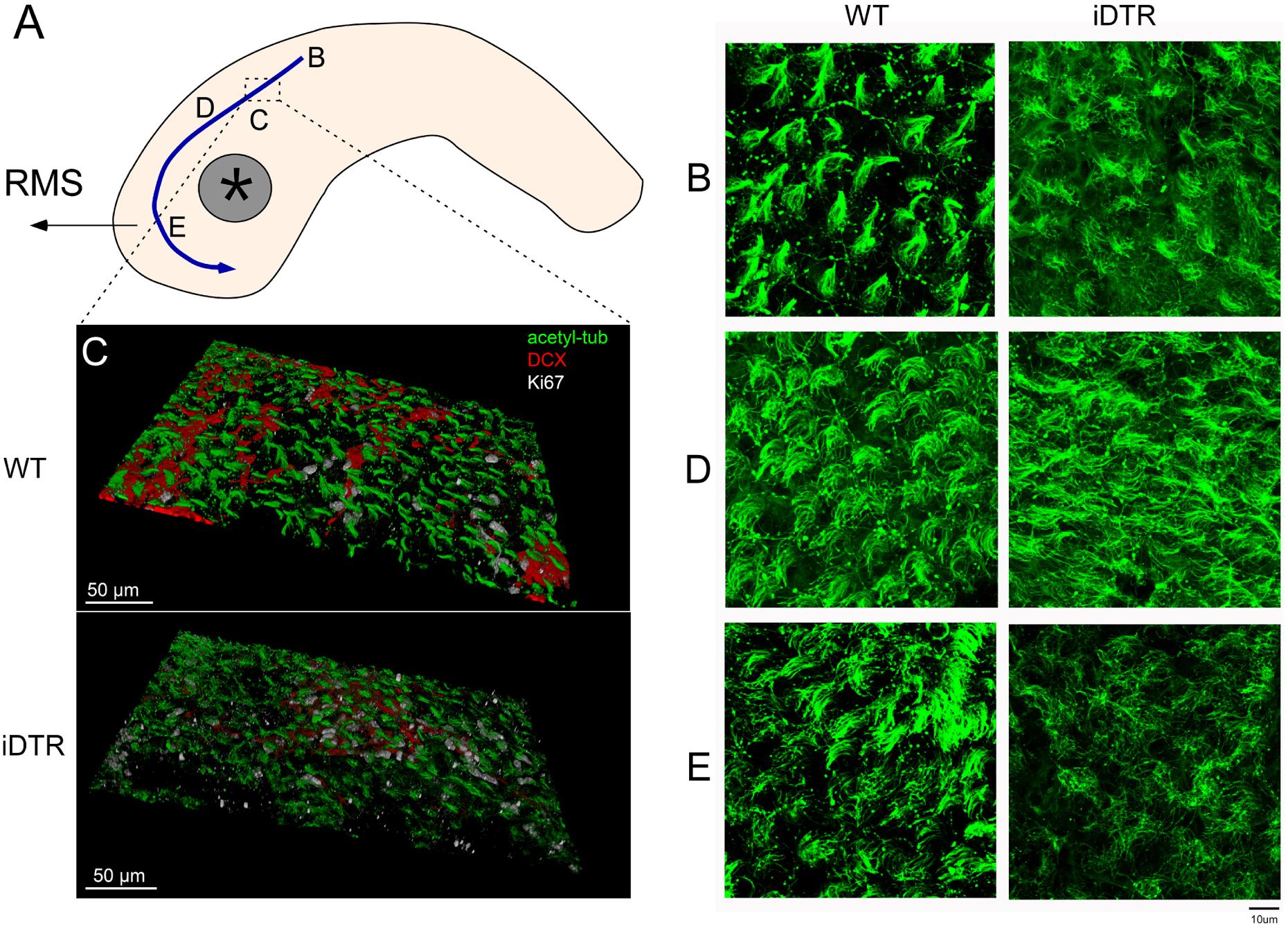
Disrupted ependymal cilia structure in ChP-ablated iDTR mice. (A) – schematic of the SVZ wall with the path of ependymal flow (blue arrow) and the attachment point (grey ceircle with asterisk). RMS – rostral migratory stream. B, D, E – representative areas in the ependymal flow path (A) with distinct directional cilia bundles at 3 months post-Dtx, 6 month-old animals. Stained for acetylated tubulin (green). Scale bar – 10 µm (C) – representative 3D reconstructions of the ventricular surface stained for acetylated tubulin (green), DCX (red) and ki67 (gray). Scale bar – 50 µm.

#### The choroid plexus/CSF are essential for newly-born neuroblast retention in the SVZ

While it has previously been hypothesized that the CSF and CSF-borne factors are essential for adult neurogenesis and particularly for NSC proliferation in the SVZ^16,20^, our results suggest that the presence of CSF flow or ChP-derived factors could instead be critical for maintaining the ventricular wall localization of DCX+ newly-born neuroblasts. In order to further examine the fate of newly-born neuroblasts after ChP ablation, we performed 5-bromo-2’-deoxyuridine (BrdU) labeling (2 injections 4 hours apart) at 5 months after Dtx treatment and harvested tissue at 1 week and 1 month post-labeling (Fig. 8 A). To evaluate whether the homeostatic continuous migration and addition of newly-born neuroblasts from the SVZ into the OB is affected, we performed BrdU immunostaining in sagittal brain sections (Fig. 8). As expected, at 1 week post-labeling, BrdU+ cells were enriched predominantly in the rostral migratory stream (RMS) compared to the granular cell layer (GCL), while at 1 month they have largely transitioned into the GCL from the RMS. This temporal transition pattern was observed in both WT and ChP-ablated mice. However, in ChP-ablated animals, we observed a pronounced increase in BrdU-labelled cells in the RMS at 1 week post-labeling compared to WT mice (Fig. 8 B, C, F – I). At one-month post-labelling, this increase was sustained in ChP-ablated mice in the GCL, with the majority of BrdU+ cells being NeuN+ granule neurons (Figs. 8 D, E, J – M). We also performed BrdU labeling using a different dosage regimen to capture multiple cell division cycles (1 injection per day for 3 consecutive days) at 6 months after ChP ablation and collected OB sections in the coronal plane at 1 month post-labeling (Supplementary Fig. 6). In this labeling paradigm in an independent cohort of mice, we observed a similar phenotype of increased BrdU+NeuN+ cells in the GCL of ChP-ablated mice at 1 month post-BrdU pulse labeling.

**Figure 8.**
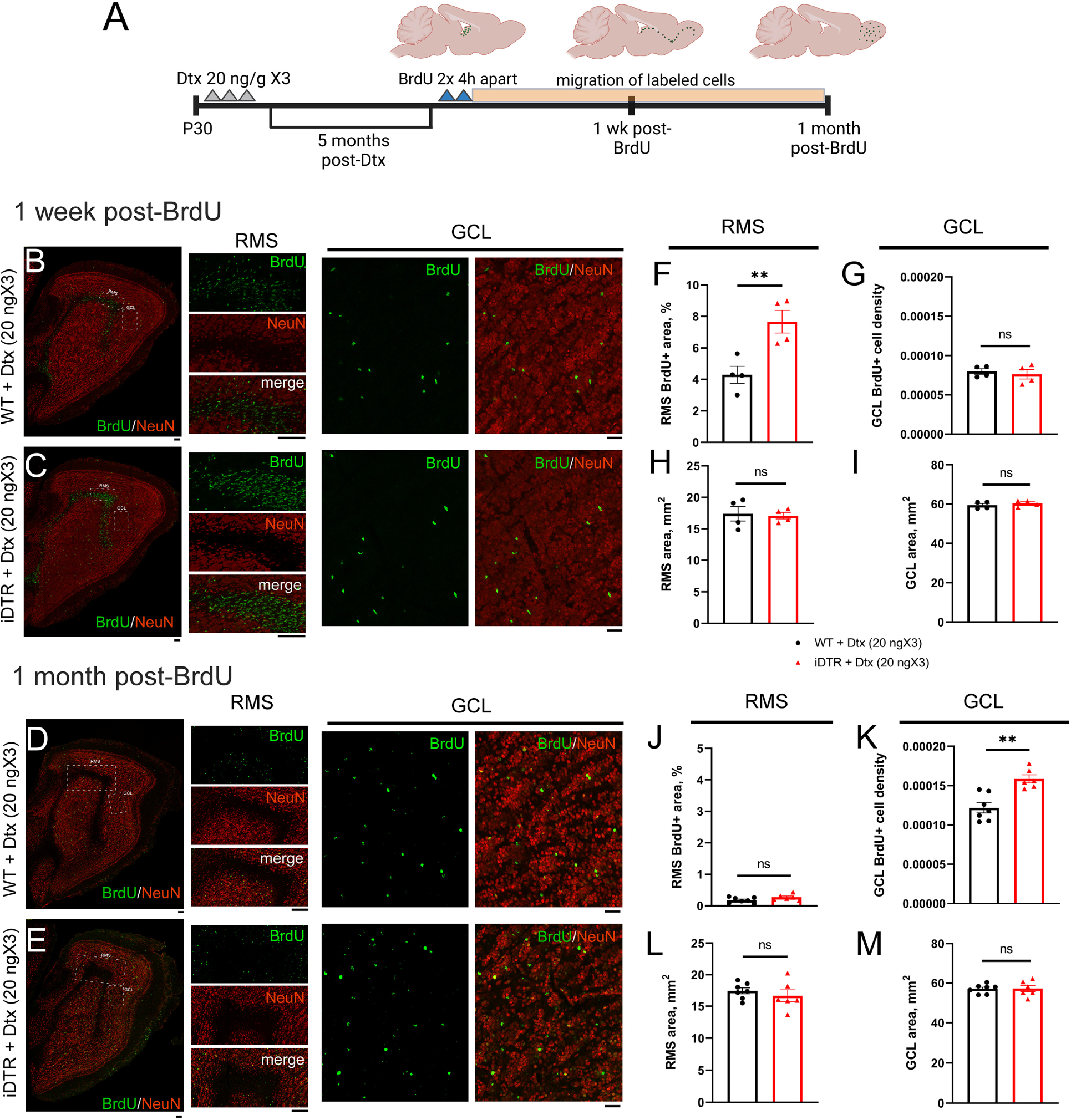
Effects of ChP ablation on migration of newly-born neuroblasts into the OB. A – experimental design schematic. BrdU labeling was performed at 5 months post-Dtx and tissue was collected at 1 week and 1 month post-BrdU. Representative OB sagittal sections from Dtx-treated WT (B, D) and iDTR (C, E) animals at 1 week (B, C) and 1 month (D, E) post-labeling, stained for BrdU (green) and NeuN (red), with magnified portions showing representative areas of the RMS and GCL. (F, H, J, L) – quantification of RMS BrdU+ area fraction (F, J) and RMS area (H, L) at 1 week post-BrdU (F, H) or 1 month post-BrdU (J, L). (G, I, K, M) – quantification of GCL BrdU+ cell density (G, K) and GCL area (I, M) at 1 week post-BrdU (G, I) or 1 month post-BrdU (K, M). Scale bar – 100 µm. Data are mean ± SEM, n = 4 per group or 1 week (F – I), n = 6-7 per group for 1 month (J – M), 3-4 sections per animal. Student’s t-test or Mann-Whitney U test. ns – p ≥ 0.05, ** – p < 0.01.

These unexpected results suggest that the decreased number of DCX+ newly-born neuroblasts in the SVZ wall after ChP ablation may not be due to decreased differentiation or survival but may instead be due to enhanced migration into the OB/RMS at 1 week after pulse labeling (Fig. 8 B, C) and subsequent maturation into BrdU+/NeuN+ adult-born neurons at 1 month later (Fig. 8 D, E). Thus, the role of the ChP and the CSF in adult SVZ neurogenesis could be to retain the newly-born neuroblasts in the SVZ and delay their migration into the OB. Indeed, using whole mount SVZ immunostaining, at 1 week post-labeling, we observed a corresponding pronounced decrease in BrdU+ cell population in the SVZ wall of ChP-ablated mice (Supplementary Fig. 7 B, H), while total Ki67+ proliferating cell number was unaltered (Fig. 6 I, L, Supplementary Fig. 7 F, G), suggesting that the increase in labeled cells in the OB/RMS at 1 week post-labeling is likely not due to differences in BrdU labeling efficiency, but due to increased exit of BrdU labeled cells from the SVZ into the OB.

#### ChP ablation impairs the neurogenic response to ischemic stroke

We next hypothesize that the decreased DCX+ neuroblast retention at the SVZ wall in ChP-ablated mice could potentially affect the dynamics of adult neurogenesis and the migration of neuroblasts into the injured area in response to CNS injury such as stroke. To determine the role of the ChP and CSF in post-stroke adult neurogenesis and neuroblast migration, we performed transient middle cerebral artery occlusion (tMCAo) surgery to induce focal cerebral ischemia in cohorts of WT and iDTR mice at 3 months post-Dtx (Fig. 9 A). Occlusion was maintained for 60 minutes and then the occluding microfilament was withdrawn to induce reperfusion. Successful infarct induction was validated via MRI at 24-48 hours post-MCAo (Fig. 9 B, C). To note, no statistically significant differences in edema volume, infarct volume and percentage of edema expansion were observed between WT and ChP-ablated mice (Fig. 9 D, E, F). Serial coronal brain sections were collected at 35 days post-stoke (a time point at which both SVZ neurogenesis and longitudinal neuroblast migration have been shown to be stimulated) and stained for DCX to evaluate the neurogenic response to ischemia and neuroblast migration into the lesion site (Fig. 9 G – J). On the side ipsilateral to the stroke, we observed a marked decrease in DCX+ cells in the striatum of ChP-ablated mice compared to WT stroke mice, indicating a reduction in neuroblast migration into the lesion site (Fig. 9 M). Furthermore, DCX immunopositive area was also profoundly reduced at the SVZ wall in iDTR mice compared to WT, in both the ipsilateral (Fig. 9 K) and contralateral SVZ (Supplementary Fig. 8 C), indicating reduced stimulation of adult SVZ neurogenesis in response to stroke. We did not observe an effect of ChP ablation on DCX+ cells in the ipsilateral (Fig. 9 L) and contralateral (Supplementary Fig. 8 D) corpus callosum, a brain area that does not have direct contact with CSF, which consistent with the data from non-stroke animals (Fig. 6 C, D). These results suggest that, while corpus callosum neuroblast population might not rely on ChP/CSF, the decrease in SVZ wall neurogenesis following ChP ablation could be directly related to the depletion of signals coming from the adjacent CSF. Furthermore, we measured the maximum neuroblast migration distance towards the lesion site for each of the sections analyzed, and did not observe a difference between WT and ChP-ablated animals (Fig. 9 N), suggesting that the reduced neuroblast number in the striatum is not likely due to reduced migration of neuroblasts into the lesion site, but instead is likely caused by the decreased pool of available SVZ NBs at the SVZ wall.

**Figure 9.**
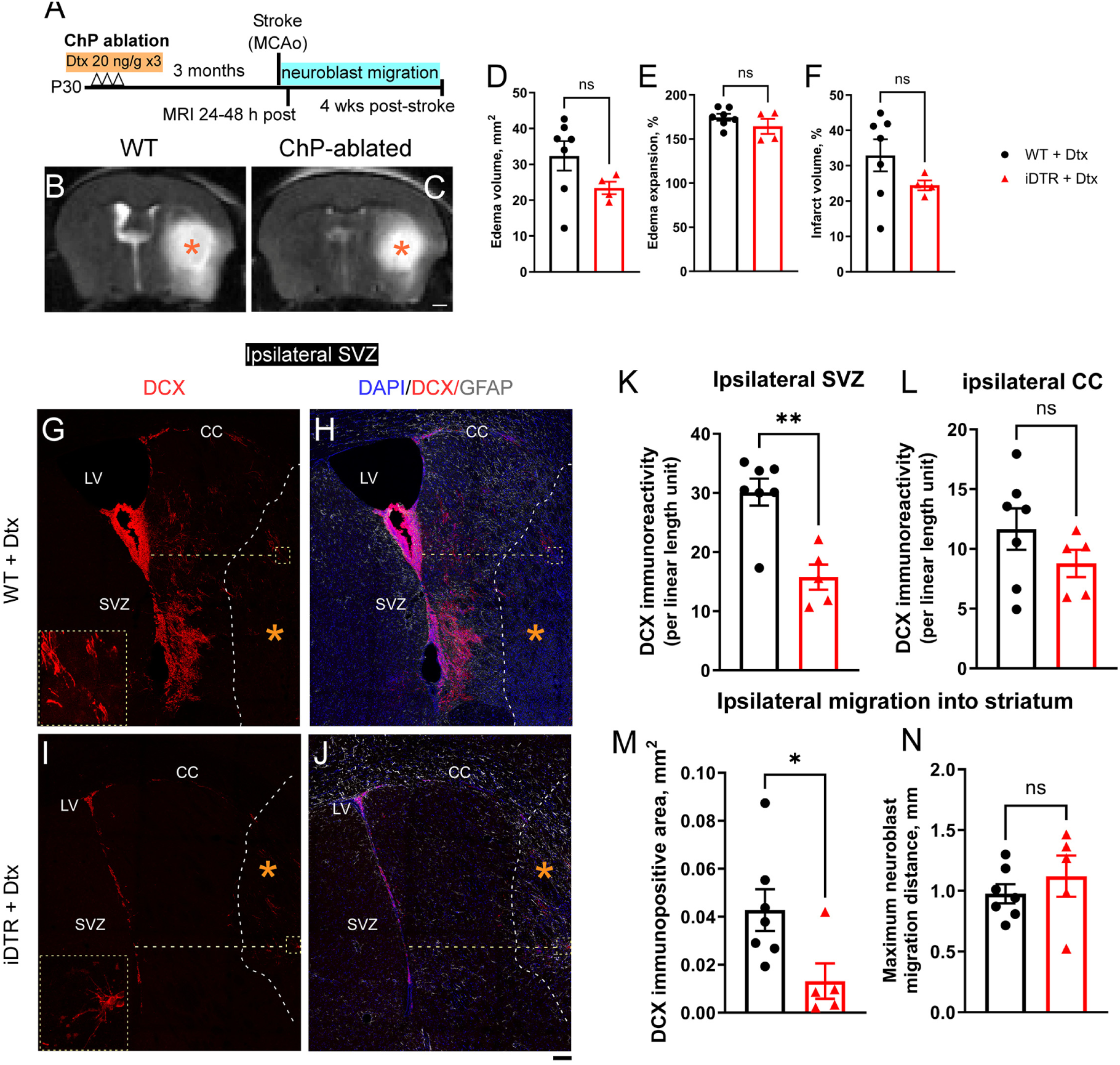
Effect of ChP ablation on SVZ neurogenesis at 1 month after ischemic stroke. A – schematic of the experimental design. (B, C) – representative T2-weighed MR images showing induction of focal cerebral ischemia in the striatum of WT (B) and iDTR (C) mice. Scale bar – 1 mm. (D, E, F) – planimetric quantification of edema volume (D), percentage of edema expansion (E), and percentage of infarct volume (F). Data are mean ± SEM, n = 4 – 6 per group, Student’s t-test, or Mann-Whitney U test. ns – p ≥ 0.05. (G-K) – representative images of coronal sections of the ipsilateral SVZ of WT (G, H) and iDTR (I, K), immunostained for DCX (G, I) and DCX/GFAP/DAPI (H, J). Dashed grey line shows lesion border. Dashed yellow line shows distance travelled by the furthest migrating neuroblast (yellow boxes and magnified inserts in panels G and I). LV – lateral ventricle, SVZ – subventricular zone, CC – corpus callosum. Scale bar – 100 µm. (K – M) – quantification of DCX immunopositive area in the ipsilateral SVZ (K), ipsilateral corpus callosum (L), normalized to SVZ or CC length, respectively, and ipsilateral striatum, as a measure of post-stroke neuroblast migration into the lesion site (M). (N) – quantification of the average maximum distance from the SVZ travelled by a neuroblast per each section. Data are mean ± SEM, n = 5-7 per group, Student’s t-test or Mann-Whitney U test. ns – p ≥ 0.05, * - p < 0.05, ** - p < 0.01.

## Discussion

### ChP epithelial cells are the main contributor of CSF production

The ChP has in general been recognized as the main contributor to CSF production, but some studies attributed only 30% of CSF volume produced to the ChP, albeit the studies were performed by invasive surgical excision of the lateral ventricle ChP in mammalian species^6,53,54^. Ependymal cells of the ventricular wall and the cerebral vascular endothelium have also been proposed as extrachoroidal CSF secretion sources^6^. However, in our study, when ChP epithelial cells are ablated in adult brain, we observe significant (>90%) loss of CSF volume, directly supporting the proposed notion that ChP epithelium is the main contributor of CSF production in adult brain or, alternatively, is required for production of CSF by extrachoroidal cells in healthy mice.

### Adult ChP epithelial cells have minimal regenerative capacity

Our results show that once the ChP is ablated in adult brain, the loss of CSF is stable and irreversible for at least 20 months, suggesting that the ChP epithelial cells have minimal regenerative capacity in the adult mouse brain, which has not been specifically tested previously. Importantly, it may carry implications for patients who underwent surgical ChP cauterization as a treatment for hydrocephalus^32^, recognizing that there may be cross-species differences. Additionally, considering that damage to the ChP has been observed in ischemic and hemorrhagic stroke^55^, as well as in traumatic brain injury^56^, our finding raises a question on the role of the ChP in long-term sequelae of these conditions. Specifically, considering that newly-born neuroblasts migrate from the SVZ into the infarct site in ischemic stroke^24,57,58^, and that their number is reduced in ChP-ablated animals (Fig. 6), it is possible that the degree of damage to the ChP in stroke could correlate with post-stroke recovery outcomes.

### ChP ablation and CSF loss does not cause acute neuronal death or an adverse motor phenotype

At day 3 after the start of Dtx treatment, apoptotic cell death was observed only in the ChP epithelium and not in other brain tissues (except for sparse apoptotic cells in the SVZ and SGZ which is also observed in the WT control mice). Surprisingly, we did not observe any overt neurobehavioral, survival, or growth differences arising from ChP ablation at one month post-ablation. We conducted a comprehensive battery of behavioral tests for motor function, circadian locomotor activity patterns, and spatial learning and working memory and did not observe any differences between WT and iDTR Dtx-treated mice at one month post-ablation (Fig. 5). This surprising finding raises questions about the extent to which the ChP and CSF are critical for CNS homeostasis in adult mice, as they have been previously considered essential for maintaining many aspects of normal CNS function^4^. It is possible that other cell types may compensate for some aspects of ChP function, even if ventricular volume cannot be maintained. The lack of mortality or an overt adverse motor or cognitive phenotype further underscores the utility of this model in long-term studies on the role of the ChP and CSF in aging or aging-associated disease models, such as Alzheimer’s disease (AD), Parkinson’s disease (PD), as well as brain injury models such as TBI, and stroke.

### ChP/CSF are not required for proliferative activity in the SVZ but are essential for retaining the newly-born neuroblast pool in the SVZ and decreasing their migration

Previously, it has been proposed that ChP-derived factors such as cytokines, hormones and growth factors are crucial for the proliferation of neural stem cells in the SVZ^20^, and other studies demonstrated that ENaC activation in response to CSF flow on the nonmotile CSF-contacting primary cilium on SVZ neural stem cells also induces proliferation through MAP kinase pathway activation^59^. However, our findings cast doubt on their necessity for NSC proliferation due to the absence of an effect of ChP ablation on Ki67+ cell number in the SVZ. Simultaneously, however, our data show that ChP ablation results in a reduced number of DCX+ newly born neuroblasts in the SVZ, which could be theoretically attributed to *1)* decreased differentiation or *2)* survival of DCX+ neuroblasts, or *3)* their enhanced exit from the SVZ and migration into the OB via the RMS. Using BrdU pulse labeling, we clearly show that ablation of ChP enhances the migration of DCX+ neuroblasts into the OB at 1 week, followed by an increase in mature adult-born BrdU+NeuN+ granule neurons in the OB at 1 month post-BrdU (Figs. 8, S6), with a corresponding reduction in BrdU+ cell number in the SVZ wall without affecting the total Ki67+ cell number (Supplementary Fig. 7). These data in sum support enhanced migration into the OB as the mechanism contributing to the reduced neuroblast number in the SVZ wall. Interestingly, previous reports suggest that the ChP or ependymal cilia may repel newly born neuroblasts from the SVZ via secreted mediators ^21^, whereas our results suggest that while the ChP/CSF are essential for ependymal cilia bundle maintenance function, ablation of the ChP enhances rather than impedes NB migration into the OB. It is possible that both pro- and anti-migration factors are present in the CSF and their balance is required for appropriate NB retention in the SVZ. However, molecular mechanisms of ChP-mediated NB retention are unknown and warrant further investigation in future studies.

### Loss of ependymal cilia polarity at the SVZ wall after ChP ablation

Considering the robust phenotype of disordered, diffuse ependymal cilia bundles that we observed in the SVZ whole-mounts of ChP-ablated mice (Fig. 7), our data suggests that the loss of planar polarity may contribute to reduced SVZ neuroblast retention after ChP ablation. Ependymal motile cilia are essential for maintaining CSF flow through the ventricular spaces. Importantly, ependymal cilia polarity directs the flow of CSF adjacent to the SVZ wall and guides the migration of newly-born neuroblasts in RMS^20,21^. A likely mechanism behind the emergence of cilia disorganization in ChP-ablated animals may include the loss of secreted factors or the hydrodynamics of CSF flow that are required for maintaining cilia organization and/or structure, or mechanical deformation of cilia bundles due to compression by the surrounding tissue after ventricle collapse.

### Impaired post-stroke neurogenesis after ChP ablation

Both in humans and rodents, focal cerebral ischemic stroke is known to elicit a pronounced neurogenic response in the SVZ^58^. In the SVZ, newly-born neuroblasts have been shown to migrate from the SVZ into the lesion site, and work by our group and others demonstrates that enhancing SVZ neurogenesis and/or neuroblast migration leads to improved recovery of motor function in rodent models^25–27^. Our results demonstrate that in ChP-ablated animals, the stimulation of SVZ (Figs. 9, S8) neurogenesis in response to stroke, as well as neuroblast migration into the lesion site (Fig. 9 M), are noticeably impaired. Our data suggests that ChP/CSF is required to retain the newly-born neuroblast pool in the SVZ, therefore, a possible protective role of the ChP/CSF in stroke could be to maintain a consistent population of newly-born NBs so that they could be available to facilitate neurorepair in case of cerebral ischemia. This potential role of the ChP in neurorepair could also be applicable to TBI and other CNS injuries, as well as aging, which are often characterized by CSF abnormalities^60^, including reduced production^42,47^ and flow^45^ accompanied by potential degenerative changes in the ChP^56,61^.

### New inducible mouse model for choroid plexus ablation

The robust phenotype of ChP epithelial cell ablation and shrinkage of ventricular spaces with loss of CSF volume in the iDTR mice after Dtx administration makes it an attractive model for studying ChP and CSF function in the adult mouse brain. The ChP and CSF are thought to play a role in multiple developmental^1,2^, homeostatic^6^ and disease processes^3^. Despite the importance and therapeutic potential of these understudied CNS compartments, the tools to efficiently manipulate the ChP/CSF in the adult brain have been largely unavailable. To date, the commonly available tools to study the CSF system include invasive infusions or injections of artificially introduced solutes, which have been demonstrated to directly affect glymphatic-mediated CSF circulation. Another tool used in the field is astrocytic AQP4 knockout to study the dynamics of perivascular solute clearance^49,62^. While this tool has produced interesting findings, it may have confounding phenotypes due to the broader role of astrocytic AQP4 in CNS physiology^63^, including affecting astrocyte volume^64^ and migration^65^, as well as modulating CNS immune responses^66^. Thus, the scope of such studies is necessarily limited to the dynamics of CSF/interstitial fluid exchange within the perivascular space and clearance of artificially introduced solutes without examining the role of ChP beyond CSF production, such as the functions of its extensive secretome^8^. While tools to manipulate the ChP, such as the Foxj1-CreERT2 line^67^ or rAAV 2/5^68^, have also been utilized in the field, these approaches have limited specificity for ChP epithelial cells ^67,69^While the Foxj1-CreERT2 line and adenoviral approaches allow for the manipulation of the ChP secretome^69,70^ or CSF absorption in hydrocephalus^68^, these methods do not readily permit direct manipulation of normal CSF production and flow, which largely limits our knowledge of the role of ChP/CSF in the adult brain. Hence, the newly identified specific expression of the DTR gene in the ChP epithelium could serve as a useful non-invasive tool that can break new ground in studies of the ChP-CSF system and glymphatic circulation in the mouse brain, especially considering the surprising lack of increased mortality, differences in body weight or effects on overt motor and cognitive functioning in ChP-ablated mice at 1 month post-Dtx (Fig. 5). The key features of the iDTR mouse model include: ***A***). Rapid and non-invasive ablation. Loss of CSF volume is apparent as early as day 3 after the start of Dtx treatment in heterozygous iDTR mice, allowing precise control of the timing of ChP/CSF ablation in a variety of genetic and non-genetic disease models. ***B***) Stable ablation. Ablation of the ChP and CSF volume loss induced by a single 3-day Dtx regimen is stable for at least 20 months post Dtx injection, making it feasible to study long-term effects of CSF function in aging and aging-associated diseases. ***C***) Dose-dependent ChP ablation and CSF volume loss. The 4 ng/gX3 or 20ng/gX3 Dtx dosage regimens used in the study produce different degrees of ChP ablation and CSF volume loss in adult mice at 1 month post ablation, allowing for more precision in modulating the severity of the phenotype. ***D***) Wide age range in which the ChP can be successfully ablated. iDTR mice both at 1 and 24 months of age show substantial loss of CSF volume after Dtx administration (Supplementary Fig. 4). This makes the model useful to study the role of the ChP and CSF throughout adulthood or only in aged mice, either in homeostasis or in combination with disease models. In summary, the iDTR mouse model has many key features to allow for precisely temporally controllable, dose-dependent and stable ChP ablation that can be easily combined with other animal models.

### A potential caveat in previous cell-type specific ablation studies utilizing the ROSA26iDTR mouse line

We unexpectedly discovered a Cre-independent robust ChP/CSF phenotypes in ROSA26iDTR mice after Dtx administration. While the mechanism behind Cre-independent DTR expression in the ChP is unknown, it can be speculated that alternative splicing (skipping the poly-adenlyation sequence in the LoxP flanked “stop” cassette) specific to the ChP likely plays a role. Since the publication of this transgenic mouse line in 2005^33^, its robust and timely Dtx-dependent ablation of the targeted cell types has been widely used in over 341 published studies^34^. It is possible that ablation of ChP could account for some of the results previously attributed to ablation of other cell types in prior studies. Notably, our results indicate that ChP ablation does not impair survival or result in overt behavioral deficits in several commonly used tests. Importantly, our study emphasizes the critical need for including proper experimental controls (i.e. Cre-negative iDTR+ mice + Dtx) in subsequent studies to ensure that any differences in phenotype are due to ablation of Cre expressing cells. Despite the wide application of this line, brain MRI was not utilized in prior studies, likely explaining the lack of any earlier report on this phenotype.

### Limitations

The ROSA26iDTR mouse line, while being a first-of-it-kind ChP ablation model, still possesses certain limitations, including considerations when combining this model with mouse Cre lines. As this line already contains the floxed stop cassette preceding the iDTR gene, Dtx administration must be performed before Cre induction to prevent ablation of Cre-expressing cells. Additionally, it is often challenging to distinguish between ChP-specific effects and the effects caused by the lack of CSF volume and/flow.

### Future directions

Given the growing understanding of the role of infiltrating peripheral immune cells in CNS pathology and homeostasis, and the proposed role of the ChP as the immune gateway to the CNS, this model will allow for direct testing of the proposed role of the ChP as a crucial regulator of immune function in the brain^71–74^. Our data demonstrate that we identified a powerful mouse genetic tool that will facilitate examining the role of CSF in aging, neurorepair, and brain-peripheral immune interactions in CNS diseases, such as TBI^56,60^, stroke^55^, neurodegenerative and demyelinating diseases^75^.

## Materials and Methods

### Animals

All animal experiments were performed in accordance with procedures approved by the University of Cincinnati Institutional Animal Care and Use Committee in compliance with NIH guidelines. All mice were maintained on a C57BL/6 background. Wildtype C57BL/6 (Jax #000664) and ROSA26iDTR mice (Jax # 007900) were obtained from Jackson Laboratories and bred in-house. To further validate this phenotype, we also retrospectively examined independent mouse lines that carried different Cre drivers, but all shared the iDTR allele in our own lab as well as in two independent labs (Goto lab and Crone lab). These mice lines include mGFAPcre-iDTR that contains the mGFAPcre allele (Jax #012886, Luo lab) the GLAST-CreER:iDTR (Jax #012586, Luo Lab), CX3CR1-CreER:iDTR (Jax # 021160, Luo lab), an independently maintained iDTR mouse colony in the Goto lab (Jax #007900, Goto lab), and (Chx10-Cre;iDTR, Crone lab^41^).

WT or iDTR mice were subjected to Diphtheria toxin (Dtx) treatment according to details described in Supplementary information at different ages. CSF was collected from cisterna magna as described previously^77^ and in Supplementary information. Tissue from different brain regions and other peripheral organs are harvested and subjected qRT-PCR as described in Supplementary information. For mouse stroke model, transient middle cerebral artery occlusion (tMCAo) was induced in male and female WT and iDTR mice (7 months old, 6 months post-ablation, 25–35g) by intraluminal occlusion of the left MCA for 60 min with a silicone rubber-coated monofilament (Cat.602212PK10Re and 602312PK10Re, Doccol Corporation) as previously described^76^ and in Supplementary information.

### Tissue collection for Immunohistology

Animals were anesthetized and transcardially perfused with ice-cold 0.1 M phosphate buffer (pH 7.2) and then with 4% paraformaldehyde in 0.1 M phosphate buffer (pH 7.2). The brain was dissected out and post-fixed in 4% paraformaldehyde overnight at 4 °C and sequentially dehydrated in 20% and 30% sucrose in 0.1 M phosphate buffer (pH 7.2) for cryoprotection. Preparation of choroid plexus (ChP) and SVZ whole mounts are carried out as previously described^78^ and in detail in Supplementary information. Whole mounts or coronal sections are subjected to various immunostaining protocol to detect DTR protein, DCX, Ki67, BrdU, CD31, TTR and other markers as described in details in Supplementary Information. The dosage and BrdU injection time are provided in each figure with an experimental timeline.

### T2-weighted MRI and brain ventricular measurement using T2-weighted MRI anatomical scans

WT and iDTR mice that were subjected to various dosages and different durations of Dtx treatment underwent an imaging session at 1 day before Dtx injection and different days after Dtx injection by staff that were blinded to the genotypes/treatment groups. MRI studies were carried out to detect ventricular volume as described in detail in Supplementary information.

### Image acquisition and analyses

Fluorescence images were acquired using either a Leica DM5000B microscope equipped with a motorized stage and a Hamamatsu C11440 fluorescent camera and configured to be operated through the Neurolucida 2020 software suite, Nikon Ti0E upright widefield microscope or a Leica Stellaris 8 Wide-field confocal microscope with LAS-X software. For details on unbiased sampling and quantification for each markers, see Supplemental information. For all analyses, the sum or average of data from each mouse was treated as n = 1 for statistical analysis.

For all analyses, the sum or average of data from each mouse was treated as n = 1 for statistical analysis.

### Behavioral Tests

Open Field Test, Barnes Maze, Spontaneous alternation Y-maze tests were carried out in WT or iDTR mice subjected to Dtx treatment as described in detail in Supplementary information. Experimental timelines were provided in each figure.

### Statistics

Results are expressed as mean ± SEM. Statistical analyses were performed in Sigmaplot 12.0 and GraphPad Prism 10 using Student’s t test, and one- or two-way analysis of variance (ANOVA), as appropriate, with Tukey, Dunn’s, or Holm-Šidák post hoc tests. When the data did not meet the assumptions of normality or equal variance, the corresponding nonparametric tests were used. The significance level was set at α = 0.05. Number of animals used in each experiment plotted as single data point and are indicated in Figure legends. Graphs were made in GraphPad Prism and portions of figures were created with BioRender.com. The notation for p-values was as follows: n.s – p ≥ 0.05, * – p < 0.05, ** – p < 0.01, *** – p < 0.001, **** – p < 0.0001

## Acknowledgments

We would like to thank Dr. Rajiv R. Ratan (Weill Cornell Medicine) for his invaluable feedback on the manuscript. We thank Chet Closson and the University of Cincinnati live imaging core (supported by NIH S10OD030402) for technical support.

## Funding

This study was funded by: NIH grants 1R01AG083164 (Y.L), R01NS125074 (Y.L.), R01NS107365 (Y.L.), R21NS127177 (Y.L. & J.G.), 1F31NS125930 (A.B.), R01NS112255 (S.A.C), Hydrocephalus Association Innovator Award (J.G.), The Center for Clinical and Translational Science and Training at the University of Cincinnati funded by the National Institutes of Health (NIH) Clinical and Translational Service Award (CTSA) program, award 2UL1TR001425-05A1 (J.G.), Rudi Schulte Research Institute (J.G.), Japan Society for the Promotion of Science (E.I.).

## Author contributions

Y.L. conceptualized the study and secured funding. Y.L. and A.T. designed experiments. A.T. performed most of the experiments, acquired and analyzed data with help from A.B. (immunostaining, quantification, some behavior tests), E.I. (TUNEL staining and coronal TTR quantification) and S.B. E.M.F, J.L. acquired and processed MRI data. A.T., F.B., A.B. analyzed MRI data. S.A.C. and J.G. provided samples to independently validate the phenotype and provided critical feedback. Y.L. and A.T. drafted the manuscript. All authors read, edited, and approved the final version of the manuscript.

## Competing interests

The authors declare they have no competing interests.

## Supplementary Information

### Supplementary Materials and Methods

#### Animals

All animal experiments were performed in accordance with procedures approved by the University of Cincinnati Institutional Animal Care and Use Committee in compliance with NIH guidelines. All mice were maintained on a C57BL/6 background. Wildtype C57BL/6 (Jax #000664) and ROSA26iDTR mice (Jax # 007900) were obtained from Jackson Laboratories and bred in-house. The mice were housed in the animal facility of University of Cincinnati on a 14-h light/10-h dark diurnal cycle. Food and water were provided *ad libitum*. For PCR primer sequences used for iDTR genotyping, please see Supplementary Table 1. Both female and male adult mice were used in this study. The age of mice in each experiment is indicated in the individual figure legends. To further validate this phenotype, we also retrospectively examined independent mouse lines that carried different Cre drivers, but all shared the iDTR allele in our own lab as well as in two independent labs (Goto lab and Crone lab). These mice lines include mGFAPcre-iDTR that contains the mGFAPcre allele (Jax #012886, Luo lab) the GLAST-CreER:iDTR (Jax #012586, Luo Lab), CX3CR1-CreER:iDTR (Jax # 021160, Luo lab), an independently maintained iDTR mouse colony in the Goto lab (Jax #007900, Goto lab), and (Chx10-Cre;iDTR, Crone lab^41^).

#### Administration of Diphtheria toxin (Dtx) to ablate the choroid plexus (ChP) in the iDTR mice

Recombinant *Corynebacterium diphtheriae* toxin was obtained from Fisher (List Biological Laboratories, Cat. #150) or Sigma (cat. # D0564-1MG) as a lyophilized powder and reconstituted in 0.9% saline at 2 mg/ml. The resulting stock solution was stored at -80 °C and diluted with ice-cold saline as needed to a final concentration of 2 ng/μl. The resulting working solution was kept on ice and used within one hour after preparation. As described previously^36^, Dtx was administered i.p. for three consecutive days with no more than 24h between injections. Dtx from Fisher or Sigma generated similar results in our studies.

### Murine model of transient focal ischemia

Transient middle cerebral artery occlusion (tMCAo) was induced in male and female WT and iDTR mice (7 months old, 6 months post-ablation, 25–35g) by intraluminal occlusion of the left MCA for 60 min with a silicone rubber-coated monofilament (Cat.602212PK10Re and 602312PK10Re, Doccol Corporation). Briefly, mice were anesthetized with isoflurane. Body temperature was monitored and maintained at 37 ± 0.5 C by a homeothermic blanket control unit (Harvard apparatus). A midline neck incision was made to isolate the left common carotid artery (CCA), external carotid artery (ECA), and internal carotid artery (ICA). A silicone rubber-coated monofilament was introduced via the arteriotomy in the ECA and advanced slowly through the ICA toward the origin of the MCA according to Longa’s method^76^. To ensure consistent and successful blockage of the MCA, regional cerebral blood flow was monitored in all stroke animals by Laser Doppler flowmetry (PeriFlux system 5000, Perimed, Sweden). The filament was left in the ICA for 60 minutes, followed by withdrawal. After incision closure, mice were subcutaneously given 1mL of warm saline and placed in a heated animal intensive care unit until recovery. T2-weighted MRI imaging was performed at 24-48 hours post-stroke.

### Collection of CSF from the cisterna magna

Animals were anesthetized and the head was fixed in a stereotactic frame. As described previously^77^, the skin and the underlying muscle layers were separated to expose the atlanto-occipital membrane. A glass pipette was used to collect CSF over 12 minutes after which animals were euthanized. Next, the CSF was weighed, and the volume of the CSF was estimated by the weight of collected CSF (assumed to be approximately equal to that of water).

#### Examination of DTR expression in ChP and other tissues via qRT-PCR

To examine DTR expression in the ChP and other tissues in WT and iDTR mice, ChP, various brain regions and peripheral tissues were harvested from adult WT, homo- or heterozygous iDTR mice for quantitative RT-PCR. Mice were perfused with ice-cold phosphate buffer (pH 7.2) before being euthanized, and the brains were immediately removed and chilled on ice. The 3^rd^, 4^th,^ and lateral ventricle ChP tissues, cortical, hippocampal, thalamus and cerebellar tissues were dissected, and total RNA was extracted following the instructions from the manufacturer (RNAqueous Micro or Mini RNA extraction kit, Thermo Fisher). Additionally, peripheral epithelial tissues (skin from the ear, small intestine, and lung) were also collected for total RNA extraction. Total RNA (80-300 ng) was treated with RQ-1 RNase-free Dnase I and reverse transcribed into cDNA using random hexamers by Superscript III reverse transcriptase (Life Sciences). cDNA levels for HPRT1 (hypoxanthine phosphoribosyl transferase 1), were determined using a specific universal probe Library primer probe set (Roche), and for DTR using a TaqMan gene expression assay (ThermoFisher, 4351372, assay Rh02840489_m1) by quantitative RT-PCR using a Roche Light Cycler II 480. Relative expression levels of DTR were calculated using the Δ Ct method compared to HPRT1 as a reference gene for each individual sample. Comparative DTR expression between tissues was calculated using the Δ Δ Ct method and the expression was normalized to the sample with the lowest DTR mRNA levels observed in heterozygous iDTR mice (thalamus). For primers, 6-carboxyfluorescein (FAM) labeled probes and the TaqMan assay used in the quantitative RT-PCR please see Supplementary Table 1. All RNA samples were treated with DNAse before reverse transcriptase reaction, and (-) RT reactions were included as negative control to ensure the absence of genomic DNA contamination.

#### Tissue collection for Immunohistology

Animals were anesthetized and transcardially perfused with ice-cold 0.1 M phosphate buffer (pH 7.2) and then with 4% paraformaldehyde in 0.1 M phosphate buffer (pH 7.2). The brain was dissected out and post-fixed in 4% paraformaldehyde overnight at 4 °C and sequentially dehydrated in 20% and 30% sucrose in 0.1 M phosphate buffer (pH 7.2) for cryoprotection.

#### Preparation of choroid plexus (ChP) and SVZ whole mounts

After transcardial PFA perfusion, the cerebellum, pons and medulla were separated, and a sagittal cut was made to separate the hemispheres. Using fine forceps and a stab knife, the cerebral cortex was peeled away, and the midbrain and thalamus dissected, exposing the ventricle wall. The lateral ventricle ChP was isolated and collected in 0.1 M PB for further staining. The whole lateral ventricular wall containing the thin layer of SVZ was dissected to prepare for SVZ whole mount IHC as described previously^78^ (also see *SVZ whole-mount staining for characterization of neurogenesis and ependymal cell motile cilia* for details below).

#### Immunostaining

##### Free-floating IF staining of coronal sections or whole choroid plexus tissues

For free-floating immunofluorescent staining, the brain tissue was frozen in OCT on dry ice and sectioned at 30 μm on a cryostat. Whole dissected ChP tissue (collected at 1 month post-ablation) or coronal brain sections were washed 3x and blocked in blocking buffer (4% BSA, 0.3% Triton X-100 (Acros Organics) in in 0.1 M phosphate buffer, pH 7.2) for 1 hr at RT with shaking and transferred into primary antibody solution (For antibodies used please see Supplementary Table 1) in blocking buffer for incubation overnight at 4 °C on a shaker.

Then, sections were washed in PB before incubated in secondary antibodies (conjugated with Alexa 488, Alexa 555, Alexa 647, Alexa 790 1:500-1000; Life Technologies, Carlsbad, CA, USA or Jackson Immuno Research, West Grove, PA, USA) dissolved in blocking buffer for 4 hours at RT with shaking. For the ChP whole mounts, the procedure was the same except for the secondary antibody incubation being done overnight at 4 °C with shaking. Free floating sections were mounted onto microscope slides (12-550-15, Fisherbrand) and coverslipped in Mowiol 4-88 (17951, Polysciences Inc.) mounting medium.

##### Staining of directly mounted coronal cryosections

For ChP area quantification using coronal brain sections, brain tissue was sectioned coronally at 40 um into 4 sets on a cryostat and directly mounted onto glass slides to prevent potential loss of ChP tissue during free floating staining. Slides were washed 3x and blocked overnight at 4 °C in blocking buffer using CoverWell incubation chambers (645502, Grace Bio-labs), 800 ul per chamber. Then, the slides were incubated for 72 hours (to ensure good penetration) at 4 °C in primary antibody diluted in blocking buffer, washed 3×30 min and then incubated in secondary antibody diluted in blocking buffer for 48 hours at 4°C. All incubations were done in CoverWell chambers. For antibodies used please see the Supplementary Table 1.

##### TSA staining of DTR protein

For diphtheria toxin receptor (DTR) staining, tyramide signal amplification (TSA) was performed. Briefly, 40 µm-thick coronal sections were treated with 3% H_2_O_2_ solution for 1 hour, blocked, and incubated with Goat anti-HB-EGF (human) antibody (1:500, R&D Systems, AF-259-NA) overnight. On the second day, after stringent washing, the tissue was incubated with anti-goat IgG biotinylated secondary antibody for 2 hours, then ABC mix solution from VECTASTAIN ABC kit^TM^ (Vector) for 1 hour, and then tyramide dissolved in 0.1M borate buffer was applied for 10 min with stringent washing between the incubations. Streptavidin Alexa Fluor 594 (Thermo Fisher) was then applied for 1 hour, followed by DAPI (Sigma-Aldrich) staining and mounting with coverslip and DAPI Fluoromount-G mounting media (SouthernBiotech 0100-20).

##### Co-staining of TUNEL and choroid plexus cell-type markers

To determine identity of the cell types undergoing apoptosis in response to Dtx, the terminal deoxynucleotidyl transferase dUTP nick end labeling (TUNEL) assay was performed according to the manufacture’s protocol using ApopTag® Fluorescein *in situ* Apoptosis Detection Kit (Millipore). For the co-staining of TUNEL with alpha-1 subunit of Na+/K+ ATPase (α-1), TUNEL assay was performed first except the final step of applying Anti-Digoxigenin Conjugate (FITC) to avoid the heat-induced inactivation of fluorescent signals, and antigen retrieval with citrate buffer (pH 6) at 95 ℃ for 45 min was performed. Blocking for 1 hour and anti-mouse α-1 antibody (1:50, Millipore, 05-369) incubation was performed overnight. Following day, after stringent washing, the tissue was incubated with anti-mouse secondary antibody along with Anti-Digoxigenin Conjugate (FITC). For the other cell type markers (CD68 Bio Rad, MCA1957T, 1:300; PDGFRβ Cell Signaling 3169S, 1:100; Isolectin GS-IB4, AlexaFluor 568 conjugate, Invitrogen I21412, 1:100) the procedure was identical except for not including the antigen retrieval step. The sections were then washed, counterstained with DAPI for 5 minutes and cover-slipped with DAPI Fluoromount-G mounting media (SouthernBiotech #0100-20).

##### 5-bromo-2’-deoxyuridine (BrdU) immunostaining

Animals injected i.p. with BrdU labeling reagent (10 µl/g bodyweight, Invitrogen #000103) 1 week or 1 month prior were sacrificed by PB/PFA perfusion and sequentially dehydrated in sucrose solution. For Fig. 8, WT and iDTR animals were treated with 2x BrdU labeling reagent 4 hours apart at 5 months post-Dtx (20 ng X3). For Supplementary Fig. 6, WT and iDTR animals were treated with 3x BrdU labeling reagent 24 hours apart at 6 months post-Dtx (4 ng X3). Coronal or sagittal cryosections were washed 3x at RT and incubated in pre-warmed freshly made DNA denaturation solution (50% formamide, 50% 2x SSC buffer) for 2 hours at 60 °C with shaking. After being washed 3x in 2x SSC at 60 °C with shaking, the tissue was incubated in 2N HCl at 37 °C with shaking. Then, the sections were incubated 1x in borate buffer (pH 8.5) at RT with shaking for 10 min and washed 1x in PB at RT. The tissue was blocked in blocking buffer for 1 hr at RT and the standard immunofluorescent protocol was followed (see *Free floating IF staining* above).

For BrdU staining of SVZ whole mount preparations, the whole-mount immunostaining protocol was followed with an additional step of incubating the tissue in 2N HCl at 37 °C for 30 minutes before the start of the IF protocol as described previously^79^.

#### Histological verification of brain ventricle loss in various iDTR allele-containing mouse lines

The perfusion fixed brain cryosection with 30 μm in thickness were mounted (some counterstained with DAPI) and cover-slipped. The mouse brain tissue from Crone lab (Chx10-Cre;iDTR) were collected in 2015-2017, thawed, mounted and cover-slipped. The brain sections were scanned using a PathScan microslide scanner to obtain images of brain sections from + 0.5 mm to − 2.0 mm in reference to Bregma, according to a coronal atlas of the mouse brain (Franklin and Paxinos, (Franklin KBJ 1997)). The image analyzer was blinded to group and treatment information.

#### T2-weighted MRI and brain ventricular measurement using T2-weighted MRI anatomical scans

WT and iDTR mice that were subjected to various dosages and different durations of Dtx treatment underwent an imaging session at 1 day before Dtx injection and different days after Dtx injection by staff that were blinded to the genotypes/treatment groups.

MRI studies were performed on a vertical wide bore 9.4T Bruker Avance III HD scanner with a 36mm proton volume coil. Mice were anesthetized with 1.5 - 2% isoflurane and kept warm with circulating air. Temperature and respiration rate were monitored with equipment from Small Animal Instruments, Inc. (SAI, Inc., NY). T2-weighted anatomical coronal images of the brain were acquired with a fat suppressed two-dimensional (2D) rapid acquisition with relaxation enhancement (RARE) sequence^80^ using the following parameters: TR 4 sec, TE 71.5 ms, echo spacing 6.5 ms, 9 or 15 slices, slice thickness/gap 0.75/0.3 mm, RARE factor 20, receiver bandwidth 67k, averages 4, matrix 192×192, FOV 28.4×28.4 mm, and total scan time 2:24 minutes.

Ventricular volumes were quantified by outlining and measuring ventricular space for each animal from the individual T2-weighted coronal images (sections ranging from Bregma AP: +1.5 mm to AP: − 4 mm) using ImageJ. CSF/ventricular volume was calculated by multiplying the area of ventricular space by the thickness of each slice (mm) of the acquisition.

#### Planimetric quantification of ischemic stroke infarct properties from T2-weighted MRI data

T2-weighted MR images were acquired with an interval of 1.0 mm (total of 15 slices per animal) at 24-48 hr post-MCAo. Eight consecutive slices per animal were quantified using ImageJ software. ROIs encompassing the contra- and ipsilateral sides, as well as the infarct area, were traced using ImageJ. Edema volume was calculated as a sum of all ROI areas of the edema lesion. Edema expansion was calculated as 100 * (edema volume) / (∑ areas of contralateral side - ∑ areas of ipsilateral side). Infarct volume percentage was calculated as 100 * (∑ areas of contralateral side - ∑ areas of ipsilateral side) / (∑ areas of contralateral side).

#### T2 Fluid-sensitive MRI and ventricle volume quantification to characterize dose-response to Dtx and stability of CSF loss over time

To further confirm the 3D volume of each ventricle in WT or ChP-ablated mice, we developed a 3D T2 Fluid-Sensitive MRI sequence to examine CSF volume *in vivo* before and after Dtx administration.

Fluid-sensitive images of the brain were acquired with a fat suppressed three-dimensional (3D) RARE sequence using the following parameters: TR 2 sec, TE 275 ms, echo spacing 11.5 ms, RARE factor 60, receiver bandwidth 104k, averages 4, matrix 320×108×80, FOV 48×16.2×12mm, and total scan time 7 minutes 44 seconds.

WT or iDTR mice that were treated with 4ng/gX3 or 20ng/gX3 dosage regimens were subjected to the 3D Fluid-sensitive MRI scan 1-3 days before the first Dtx injection and at 5-18 days after end of Dtx treatment to examine the effects of different dosages of Dtx. To examine the stability and potential reversal of the phenotype, one cohort of mice (n=4 for control or iDTR+Dtx mice) were followed up by 3D-Fluid sensitive MRI scans for up to 20 months after Dtx injection. To examine the age range of iDTR mice in which ChP ablation can be achieved, we examined a cohort of 1-month-old iDTR mice or 24-month-old iDTR mice (Dtx dosage: 20ng/gX3) and subjected them to the same 3D-fluid sensitive MRI.

The resulting series of T2 Fluid-Sensitive images were imported into Imaris 9.8.0 and 3D ventricular surfaces were reconstructed from a series of DICOM files. The surfaces were generated using the Surfaces tool with surface detail of 0.1 and seed point diameter of 0.8. The resulting 3D ventricular surfaces were manually divided into separate ventricles where appropriate. The volumes were quantified using the Measurements tool in Imaris.

### SVZ whole-mount staining for characterization of neurogenesis and ependymal cell motile cilia

Brain tissue was harvested after 4% PFA perfusion at 1, 3 and 6 months post-ablation. SVZ whole-mounts were dissected as described^78^ and washed 3×10 min in 0.1% Triton X-100 in 0.1 M PB. Then, the mounts were blocked in 4% BSA in 0.1 M PB (pH 7.2) with 2.3% Triton X-100 for 1 hour at RT. The whole mounts were transferred into the primary antibody solution and incubated for 72 hours at 4 °C with shaking. Then, the tissue was washed 3×30 min in 0.1% Triton X-100 in 0.1 M PB and transferred into secondary antibody solution and incubated for 72 hours at 4 °C with shaking, washed 3×30 min in 0.1% Triton X-100 in 0.1 M PB, mounted on microscope slides and coverslipped.

#### Image acquisition and analyses

Fluorescence images were acquired using either a Leica DM5000B microscope equipped with a motorized stage and a Hamamatsu C11440 fluorescent camera and configured to be operated through the Neurolucida 2020 software suite, Nikon Ti0E upright widefield microscope or a Leica Stellaris 8 Wide-field confocal microscope with LAS-X software. For ChP and SVZ whole mount DCX and Ki67 quantification, tissue was scanned and automatically stitched using the Slide scanning workflow in Neurolucida. For doublecortin total immunopositive area quantification in SVZ whole mounts and coronal sections, images were analyzed in Nikon Elements AR, and the DCX immunopositive area fraction was taken to estimate the dencity of DCX+ newly-born neuroblasts in the SVZ wall. For Ki67 counts, representative coronal section confocal images were acquired and Ki67+ cells were manually counted in ImageJ 1.53, or SVZ whole mounts were scanned using the MBF SetereoInvestigator at a z-stack thickness of 40 µm to fully capture the outer layer of the ventricular wall. The scanning sites were randomly selected using an unbiased sampling grid generated in StereoInvestigator to cover 10% of the total SVZ whole mount area and Ki67+ or BrdU+ cells were counted, and the population density was estimated using the Optional Fractionator workflow in StereoInvestigator. For ChP ablation evaluation, an average of 15-20 coronal sections from 1 out of 4 sets per brain were scanned and TTR+ area for LV and 3V ChP was quantified using Nikon Elements AR. For CD31+ blood vessel length density, maximum projection images of scanned whole mount ChPs were quantified using ImageJ. Consistently located ROIs were selected based on large blood vessels as landmarks. Each blood vessel within the ROI was traced, and the total linear length of capillaries within each ROI was divided by the total ROI area to obtain the length density. For BrdU+ cell number quantification in olfactory bulb sagittal sections, 3-4 sections per animal were chosen based on consistent landmarks. Using ImageJ, the images were thresholded and converted to binary masks. Then, ROIs containing the GCL and RMS were applied and the number of BrdU+ cells was automatically calculated. For quantification of BrdU+ NeuN+ cells in coronal OB sections (Supplementary Fig. 6), a similar procedure was followed using NIS Elements. For all analyses, the sum or average of data from each mouse was treated as n = 1 for statistical analysis.

For quantification of post-stroke SVZ neurogenesis, scanned images of sections containing the ipsi- and contralateral SVZ and corpus callosum, as well as the ipsilateral striatum, including the lesion site, were acquired on the Leica DM5000B microscope. Three sections containing the most DCX+ migrating neuroblasts were chosen per animal. ROIs containing the ipsi- and contralateral SVZ, ipsi- and contralateral CC, as well as the ipsilateral striatum with the lesion site, were traced using ImageJ. The images were thresholded, and DCX+ immunopositive area was calculated. To calculate the maximum migration distance towards the lesion site, a perpendicular line was drawn from the SVZ wall to the furthest migrating neuroblast for each of the three sections per animal, and an average of the three distances was taken.

#### Behavioral Tests

##### Open Field Test

The open field apparatus (Omnitech electronics Inc, Columbus, OH) consisted of 2 sets of 8 photobeam arrays for animal horizontal activity detection and 1 set of 8 photobeam arrays for vertical activity detection. Each monitor was installed on a cube frame inside of a closed cabinet. A 42Lx42Wx31T cm Plexiglas box divided into 4 chambers to allow for simultaneous recording of two animals was placed inside of each frame. Mouse bedding was evenly placed on the bottom of the box and the feeder and water bottle were installed on the box lid. The cabinet was lit in a 12 h light/12 h dark light cycle and equipped with ventilation fans. The chambers were washed using a steam mouse cage washer between tests. Animal activity was tracked using automated Fusion software (Omnitech electronics). The locomotor activity was analyzed over a 23-hour and 72-hour periods.

##### Barnes Maze

Barnes maze (Stoelting) consisted of a gray circular platform 91 cm in diameter, at an elevation of 90 cm from the floor. On the platform surface, 20 holes of 5 cm diameter were evenly distributed around the edge. Among them, 19 out of 20 holes had a small gray square tray 2 cm in depth underneath each hole, while the remaining “escape” hole led to a 5-cm deep, 5 cm-wide, 15 cm-long escape box underneath. During the probe trial, the escape hole was replaced with the same 2 cm-deep “sham” tray. The test room was lit by LED light strips at 40 lux intensity. The testing area was encircled by black curtains. Above the center of the platform, a 500-watt light bulb and a fan were set to provide additional aversive stimulation during the test. Four colored visual cues on a white background were hung surrounding the platform to provide spatial cues during the test. Between tests, the apparatus was rinsed with warm water and wiped with 20% ethanol to eliminate olfactory cues. The test included 3 stages: Habituation, Acquisition Learning and Acquisition Probe. Habituation consisted of a single 2-minute trial per animal without visual cues or additional overhead light or fan exposure. During the Acquisition learning stage, each animal was trained for 2 trials (3 minutes duration each and 15 min inter-trial interval) for 4 days with visual cues and with the overhead light and fan on. The acquisition probe stage involved a single 3-minute trial with visual cues and with the overhead light and fan on. Animal activity was tracked in real time and analyzed using the automated ANY-Maze 7.1 behavior analysis software (Stoelting). The animals were tested at 1 month after Dtx administration.

##### Spontaneous alternation Y-maze

The Y-maze apparatus consisted of 3 identical arms angled at 120° (5 cm wide, 35 cm long) enclosed by dark plastic walls 10 cm tall. The testing area was enclosed by dark curtains and lit with LED light strips at 18 lux. The animal was placed into one of the arms and allowed to freely explore the maze for 8 minutes while its locomotor activity was recorded using an overhead-mounted camera and tracked in real time using ANY-maze 7.1. To measure spatial working memory, the alternation rate was calculated as follows: % Alternations = total number of three-arm sequences / (total entries – 2).

---

**Supplementary figure 1.**
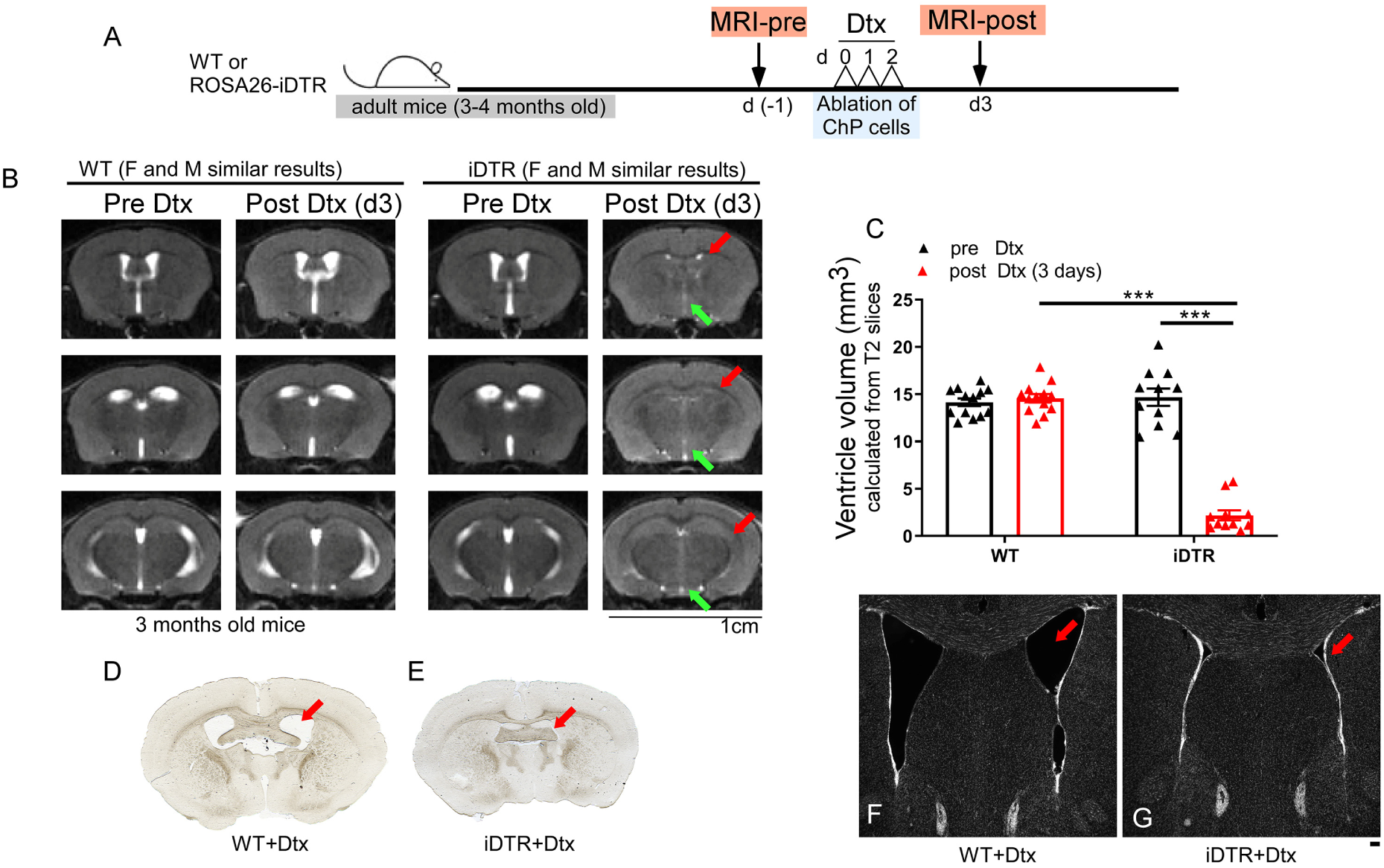
Rapid loss of ventricular spaces in Dtx-treated iDTR mice following ChP ablation. (A) Schematic of experimental timeline. (B) T2-weighed MRI one day before start of Dtx treatment and 3 days after the start of Dtx treatment, collapsed lateral and 3^rd^ ventricles shown in red and green arrows, respectively, scale bar – 1 cm. (C) Quantification of the volume of the lateral and third ventricles calculated from 9 T2 slices per animal (mean ± SEM, n = 11-13 per group, Two-way repeated measures ANOVA with genotype and timepoint as factors). (D, E) – images of coronal brain sections showing the ventricle (red arrows). (F, G) – magnified image of coronal brain sections stained with DAPI showing the ventricle lumen arrows). Scale bar – 100 µm.

**Supplementary figure 2.**
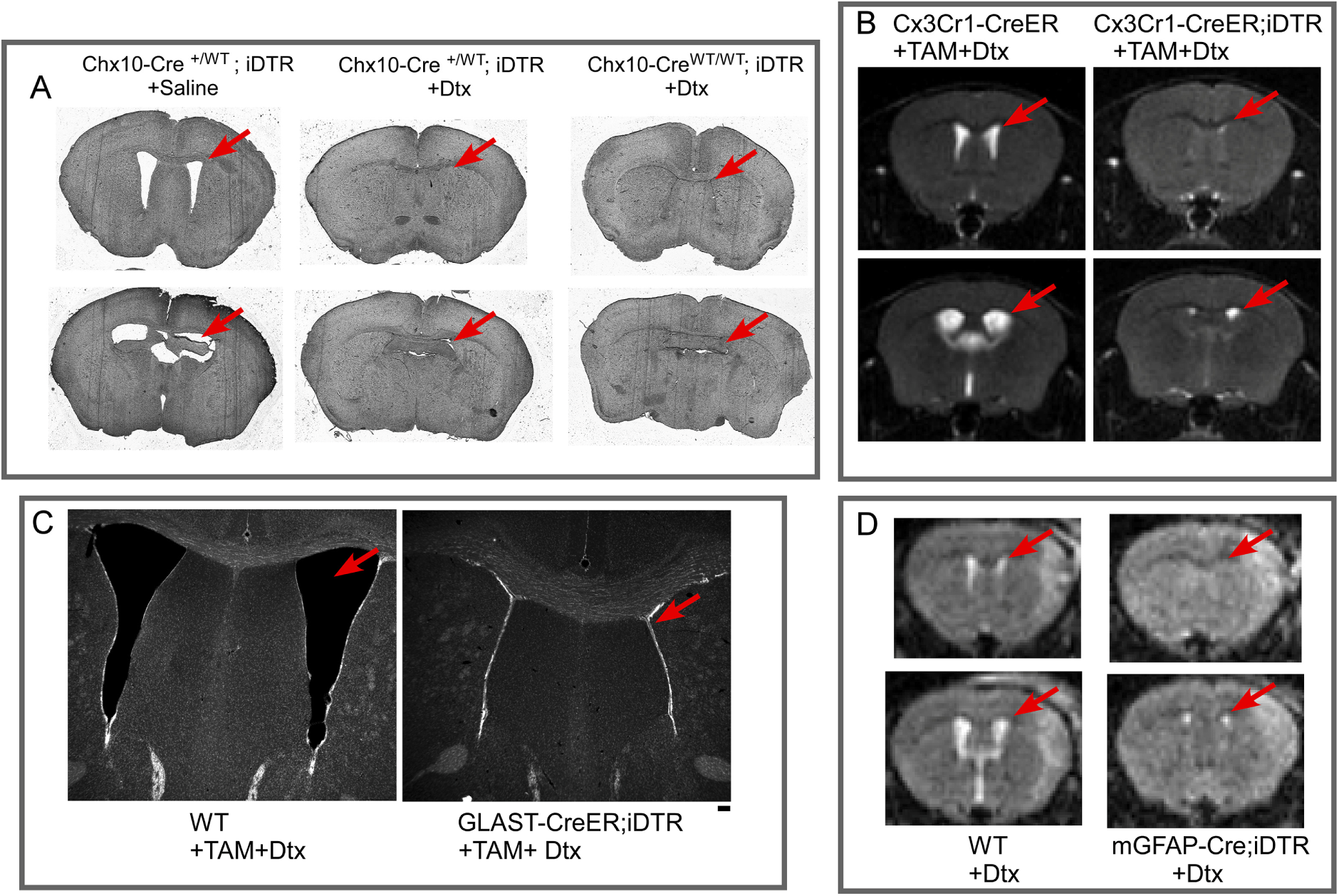
Validation of ventricular volume loss in multiple iDTR allele-containing mouse lines from different labs. (A). Histological tissue sections collected in 2015-2017 (provided by Crone lab) showing ventricle loss in Chx10-Cre+ iDTR+ and ROSA26iDTR only animals. (B, D) Phenotype of ventricular volume loss seen in T2-weighted MRI in the Cx3cr1-creER;iDTR line (expressed in macrophages and microglia) and the mGFAP-Cre;iDTR line (neural stem cells and astrocytes). (C) Histological brain sections stained with DAPI showing ventricle collapse in the GLAST-CreER;iDTR line (astrocytes), scale bar – 100 µm. In all panels the ventricle is indicated by red arrows.

**Supplementary figure 3.**
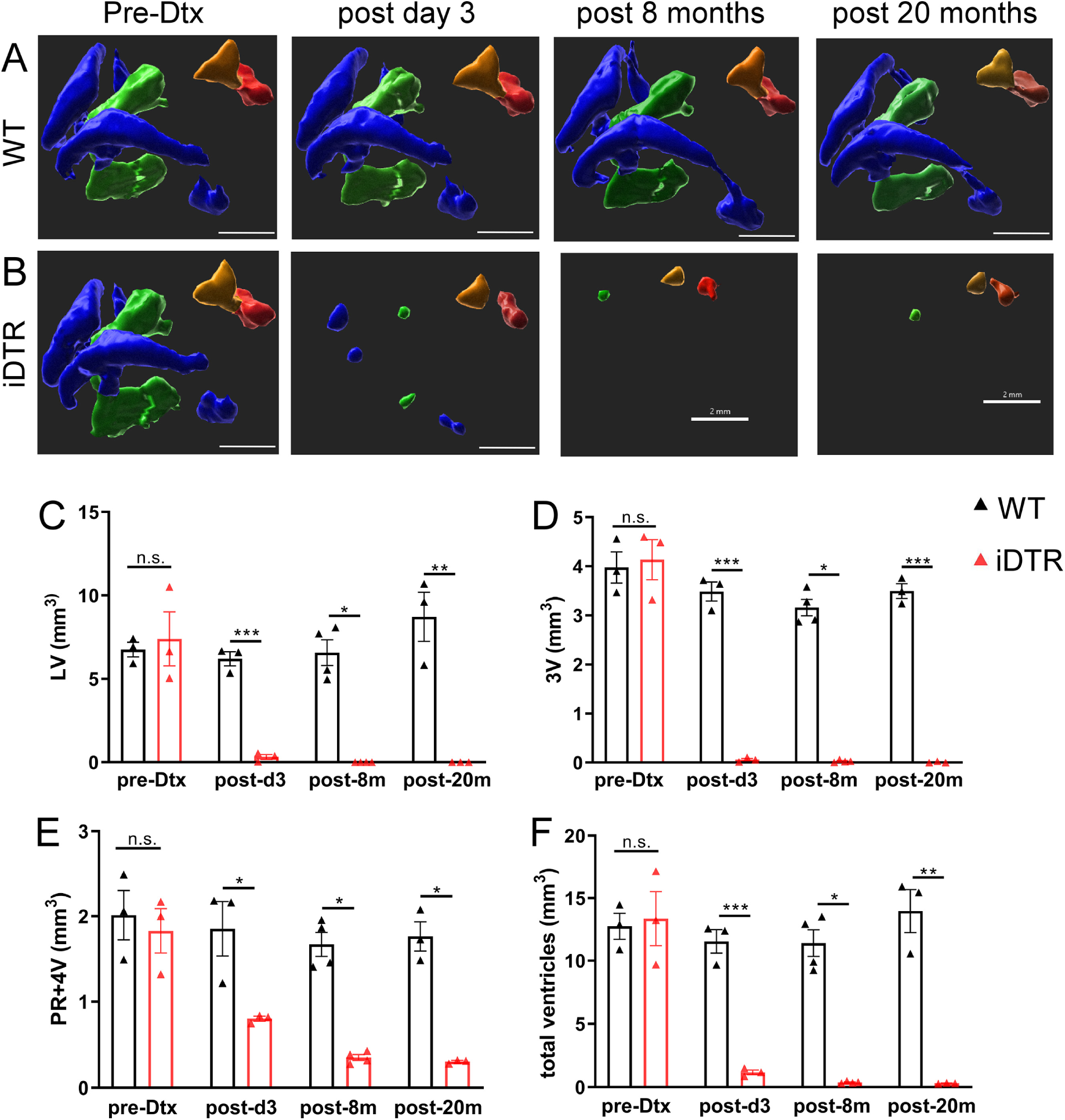
Stable and sustained CSF volume loss in Dtx-treated iDTR mice. (A, B) representative 3D reconstructions of fluid-filled ventricular spaces of WT (A) and iDTR (B) mice from T2 fluid-sensitive MRI data (lateral ventricles – blue, third ventricle – green, pineal recess – orange, fourth ventricle – red). (C-F) – quantification of ventricle volumes at day (-1) before Dtx treatment, day 3 after the start of Dtx treatment and at 8 and 20 months after the start of Dtx treatment at the age of 2 months. LV – lateral ventricles, 3V – third ventricle, PR – pineal recess, 4V – fourth ventricle (mean ± SEM, n = 3-4 per group, Student’s t-test, Welch’s t-test or Mann-Whitney U test within timepoints). n.s. – p ≥ 0.05, * - p < 0.05, ** - p < 0.01, ** - p < 0.001, *** - p < 0.0001. Scale bar – 2 mm.

**Supplementary figure 4.**
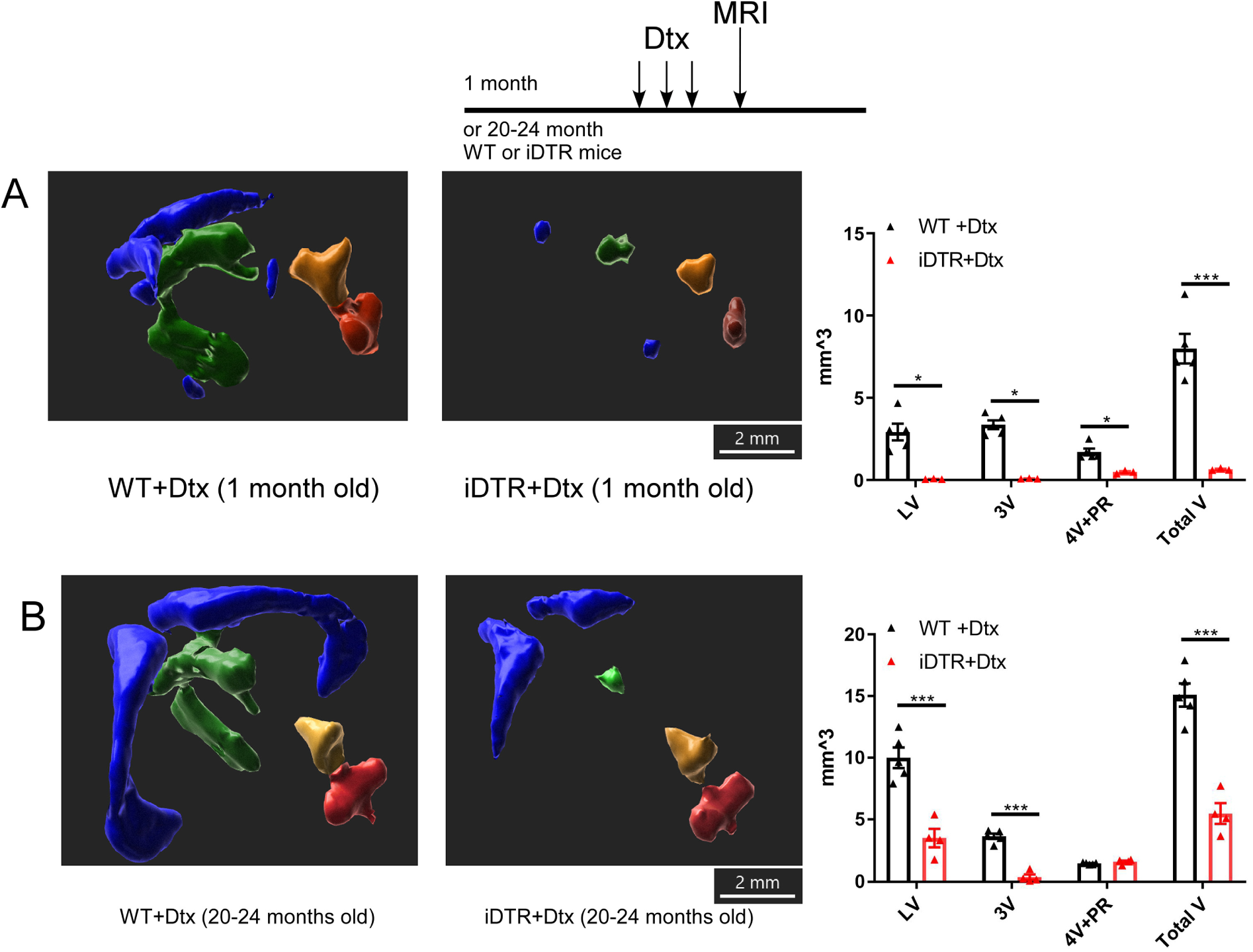
ChP ablation induces CSF volume loss in different ages across adulthood in iDTR mice treated with Dtx. (A, B) – representative 3D reconstructions and quantification of ventricular spaces of WT and iDTR mice treated with Dtx at one month of age (A) and 20-24 months of age (B), scanned at 1 month after Dtx treatment. LV – lateral ventricles (blue), 3V – third ventricle (green), PR – pineal recess (orange), 4V – fourth ventricle (red). Scale bar – 2 mm. Data are presented as mean ± SEM, n = 3-5 per group, Student’s t-test or Mann-Whitney U test. * - p < 0.05, ** - p < 0.01, *** - p < 0.001, **** - p < 0.0001.

**Supplementary Figure 5.**
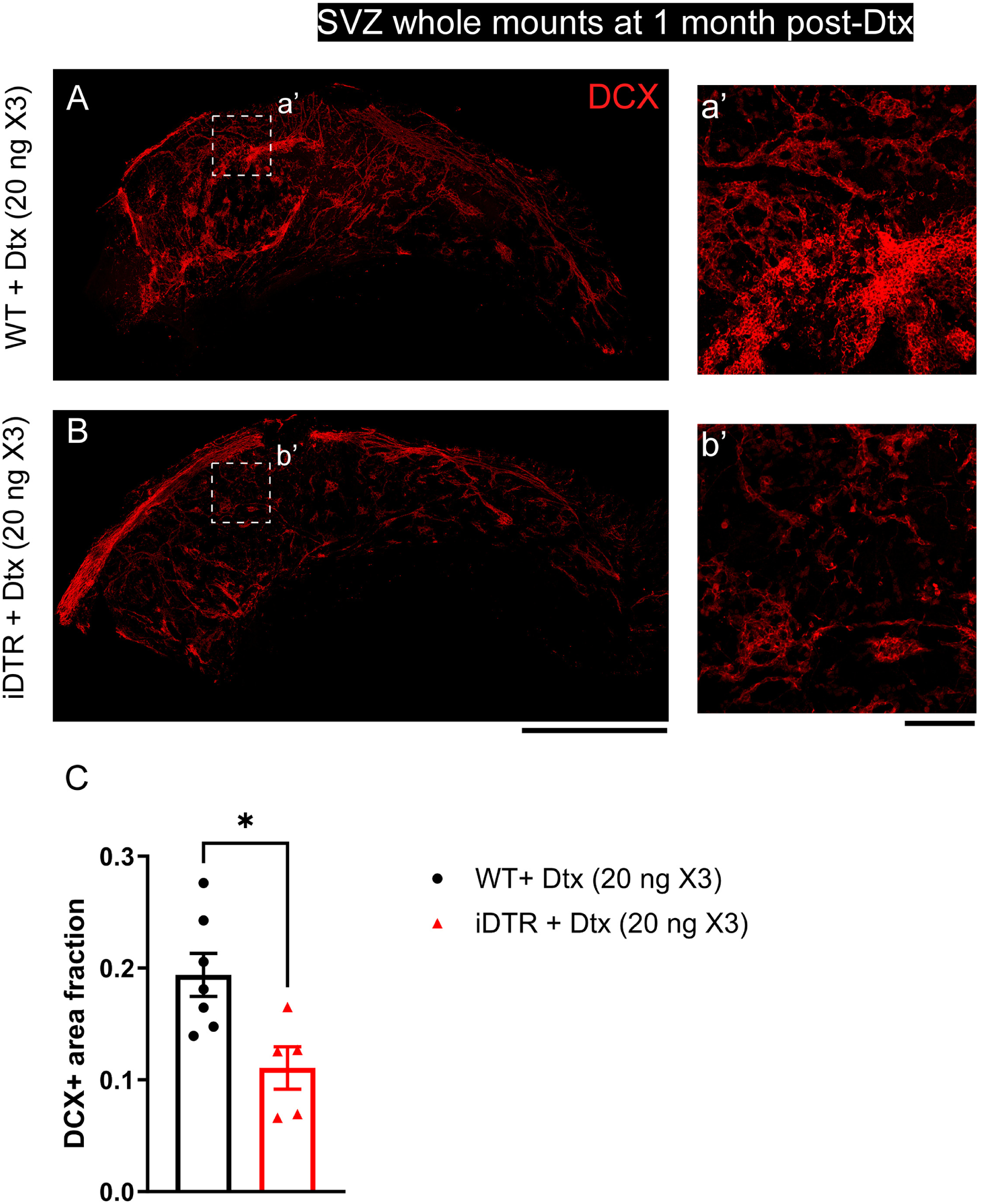
Dose-dependent effects of ChP ablation on adult SVZ newly-born neuroblast pool at 1 month post-Dtx. (A, B) – Whole-mount immunostaining for DCX at 1 month post-Dtx in WT (A) mice that received 20 ng X3 of Dtx and iDTR mice that received 20 ng X3 of Dtx (B) Scale bar – 1 mm. (a’, b’) – magnified inserts showing representative areas of the SVZ whole mounts with migrating DCX+ newly-born neuroblasts. Scale bar – 100 µm. (C) – quantification of DCX+ area fraction of the SVZ whole mounts, Student’s t-test. * - p < 0.05.

**Supplementary Figure 6.**
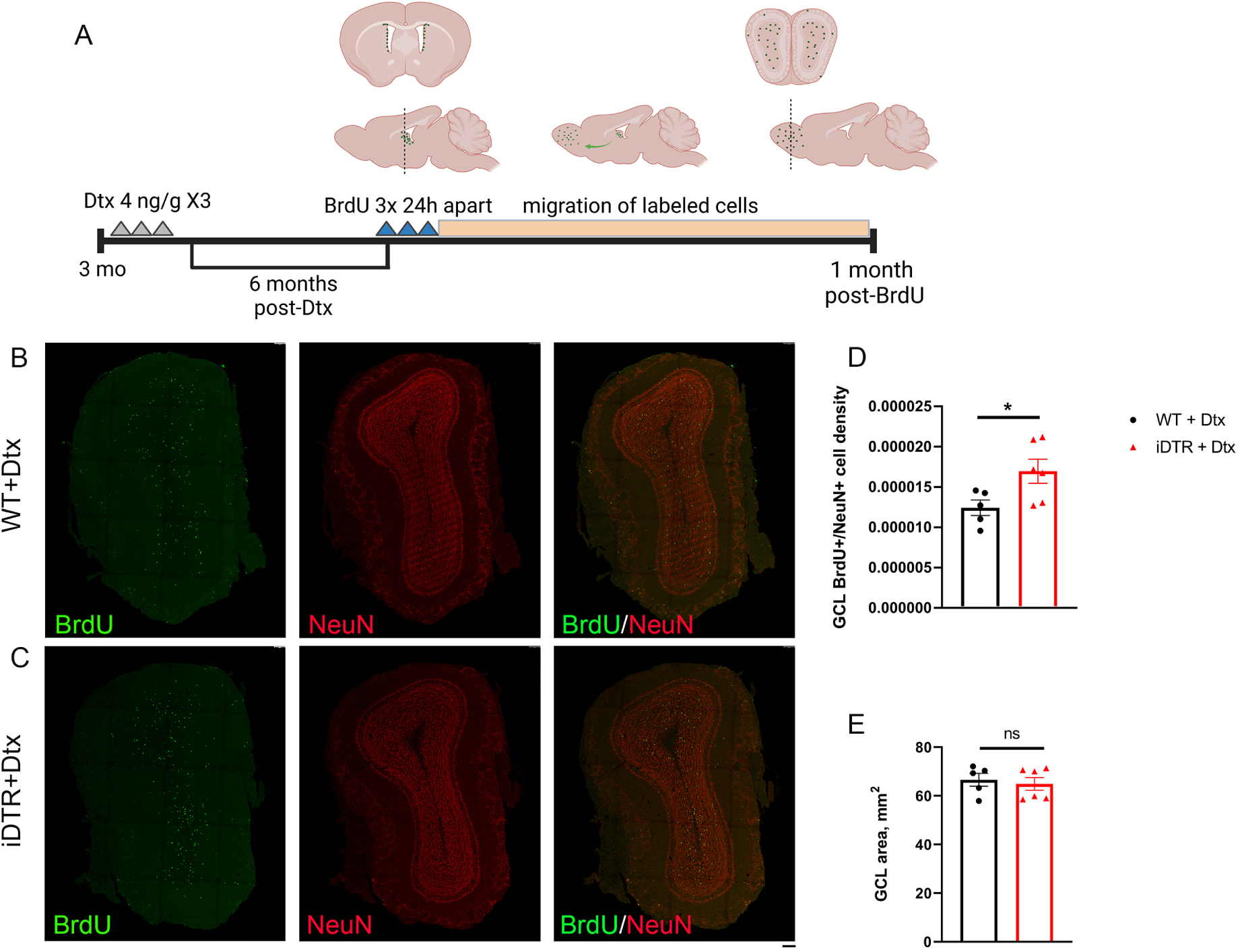
Validation of neuroblast migration phenotype using a different Dtx dosage, BrdU labeling regimen and sectioning plane (coronal). (A) – schematic of experimental design. (B, C) – representative coronal OB sections from Dtx-treated WT (B) and iDTR (C) mice stained for BrdU (green) and NeuN (red). Scale bar – 100 µm. D, E – quantification of BrdU+ cell density (D) in the GCL and GCL area (E). n = 5 – 6 per group, 4 sections per animal. Data are mean ± SEM, Student’s t-test (D) or Mann-Whitney U test (E). ns – p ≥ 0.05, * - p < 0.05.

**Supplementary figure 7.**
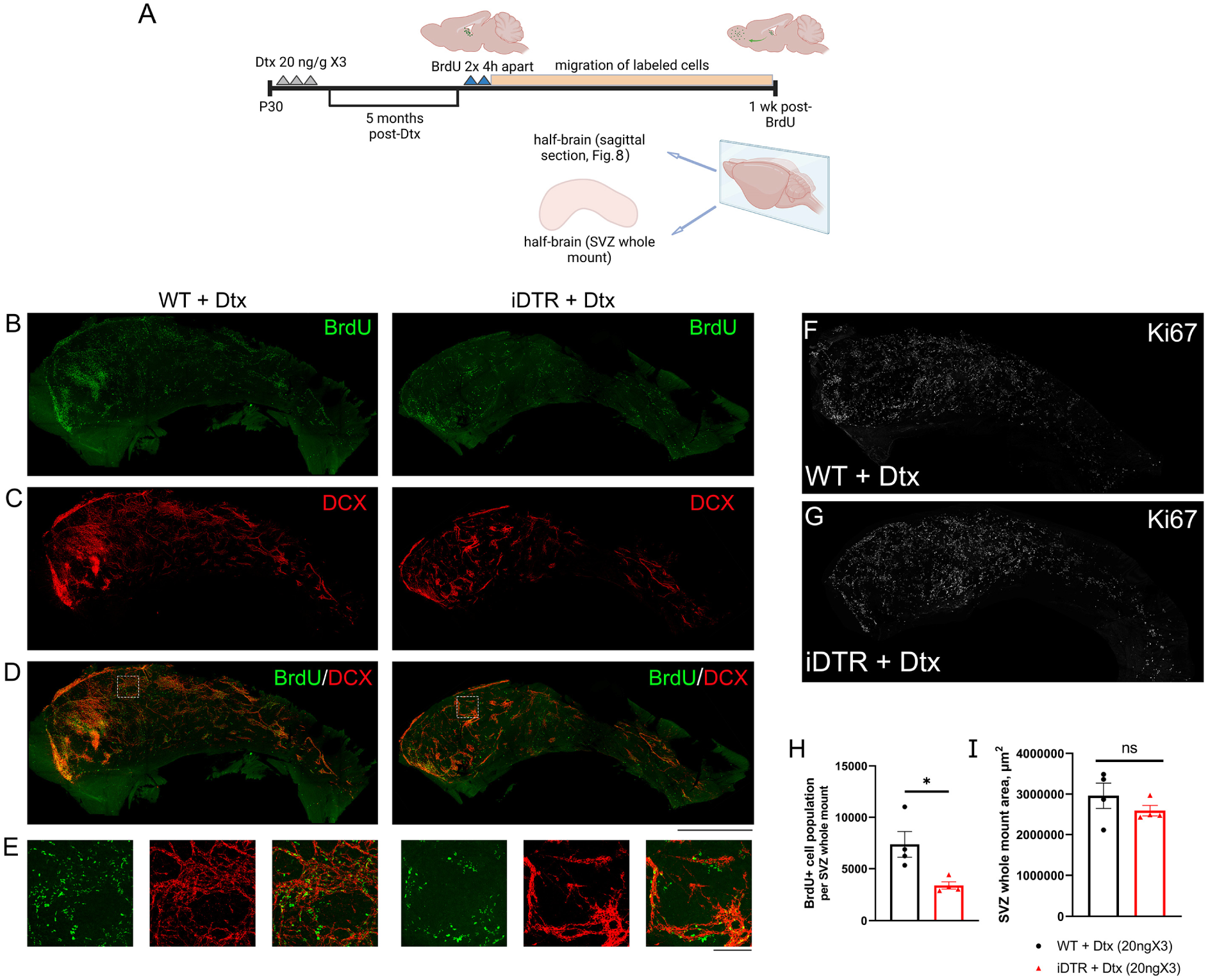
BrdU+ cell population in SVZ whole-mounts at 1 week post labeling. A – experimental design schematic. Representative images of SVZ whole mounts stained for BrdU (B) and DCX (C) Scale bar – 1000 µm. D – overlay. E – representative magnified area, scale bar – 100 µm. (F, G) – Representative images of SVZ whole mounts at 6 months post-Dtx showing similar Ki67+ cells in Dtx-treated WT (F) and iDTR (G) mice (quantification of Ki67 cells is presented in Fig. 6I and 6L). H – unbiased stereological quantification of BrdU+ cell population in SVZ whole mounts. I – quantification of SVZ whole mount area. Data are mean ± SEM, n = 4 per group. Student’s t-test. ns, p ≥ 0.05, * p < 0.05

**Supplementary Figure 8.**
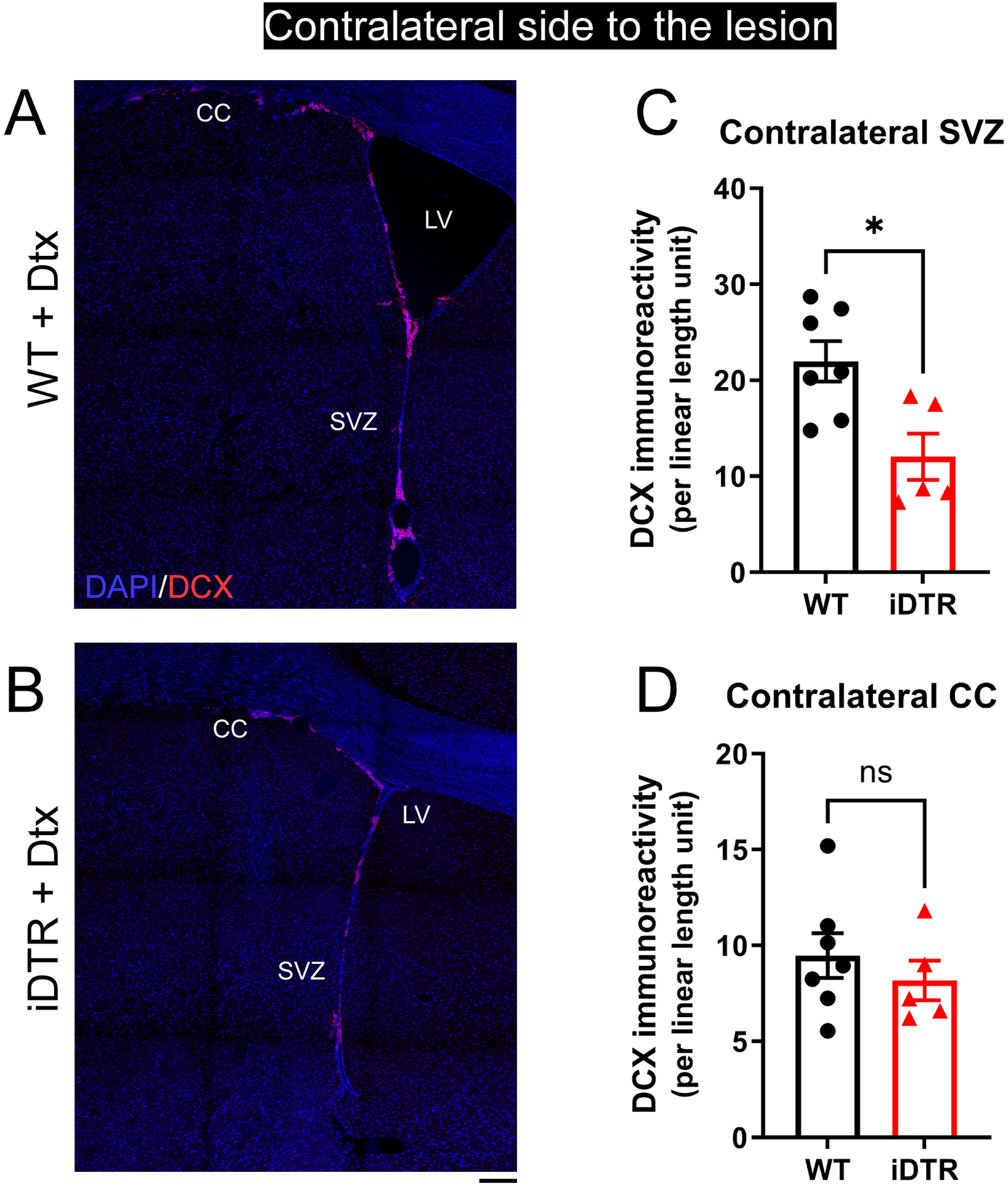
Effect of ChP ablation on the stimulation of SVZ neurogenesis in the contralateral SVZ at 1 month post-stroke. (A, B) – representative images of the contralateral SVZ of WT (A) and iDTR (B) mice, stained for DCX (red) and DAPI (blue). LV – lateral ventricle, CC – corpus callosum, SVZ – subventricular zone. Scale bar – 100 µm. (C, D) – quantification of DCX immunopositive area in the contralateral SVZ (C) and contralateral corpus callosum (D), normalized to SVZ or CC length, respectively. Data are mean ± SEM, n = 5-7 per group, Student’s t-test. ns – p ≥ 0.05, * – p < 0.05.

**Supplementary table 1.**
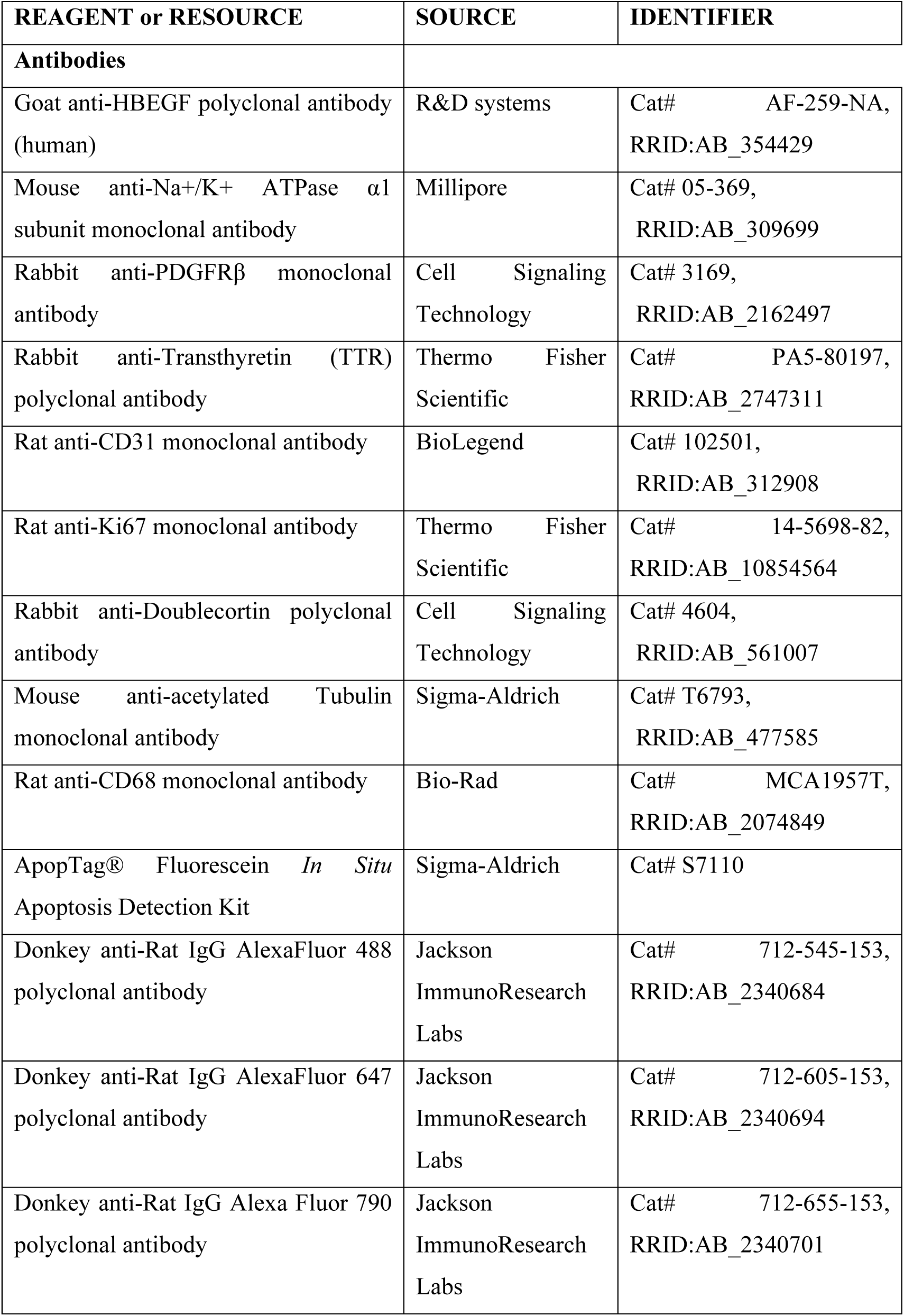
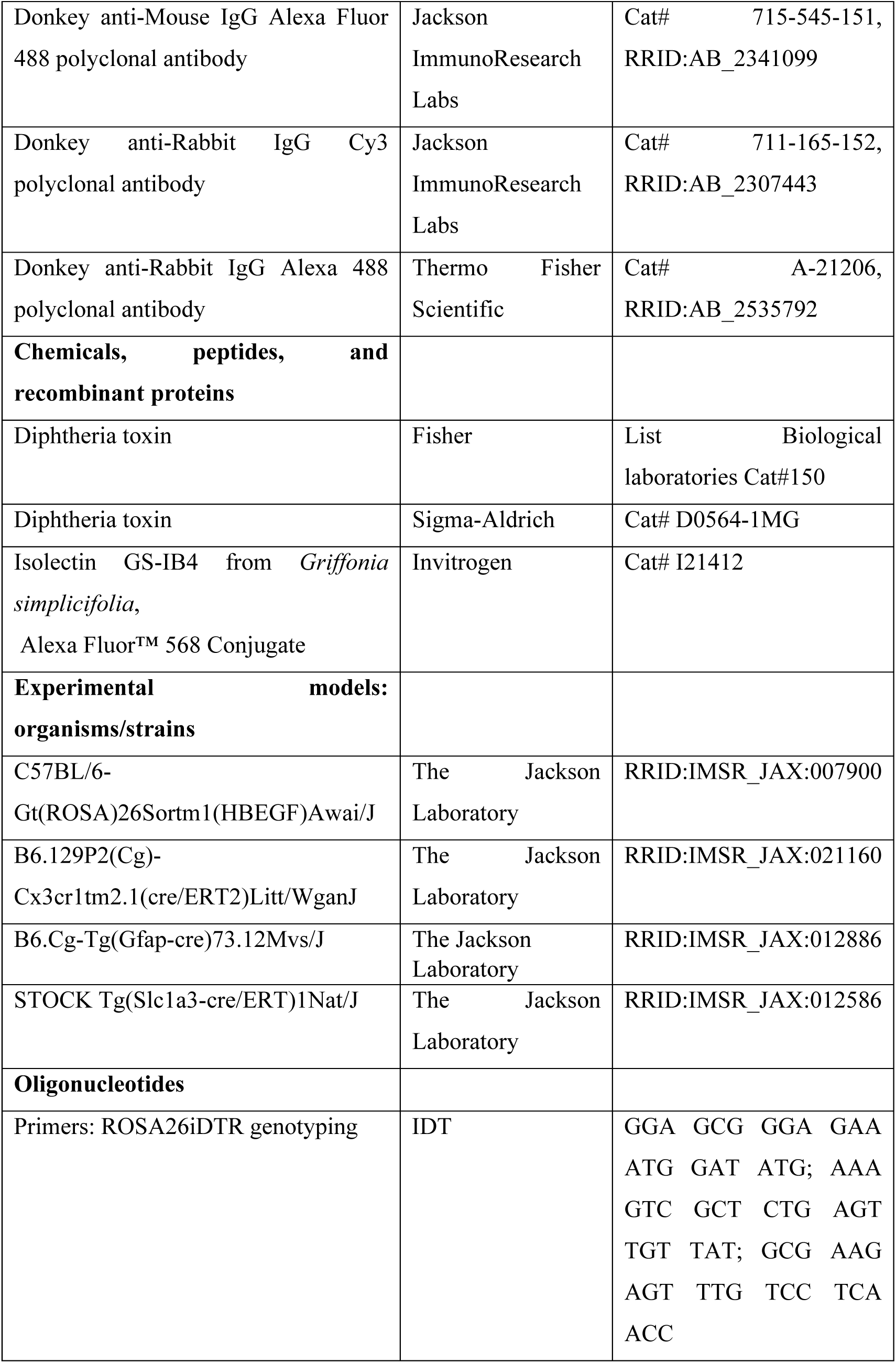
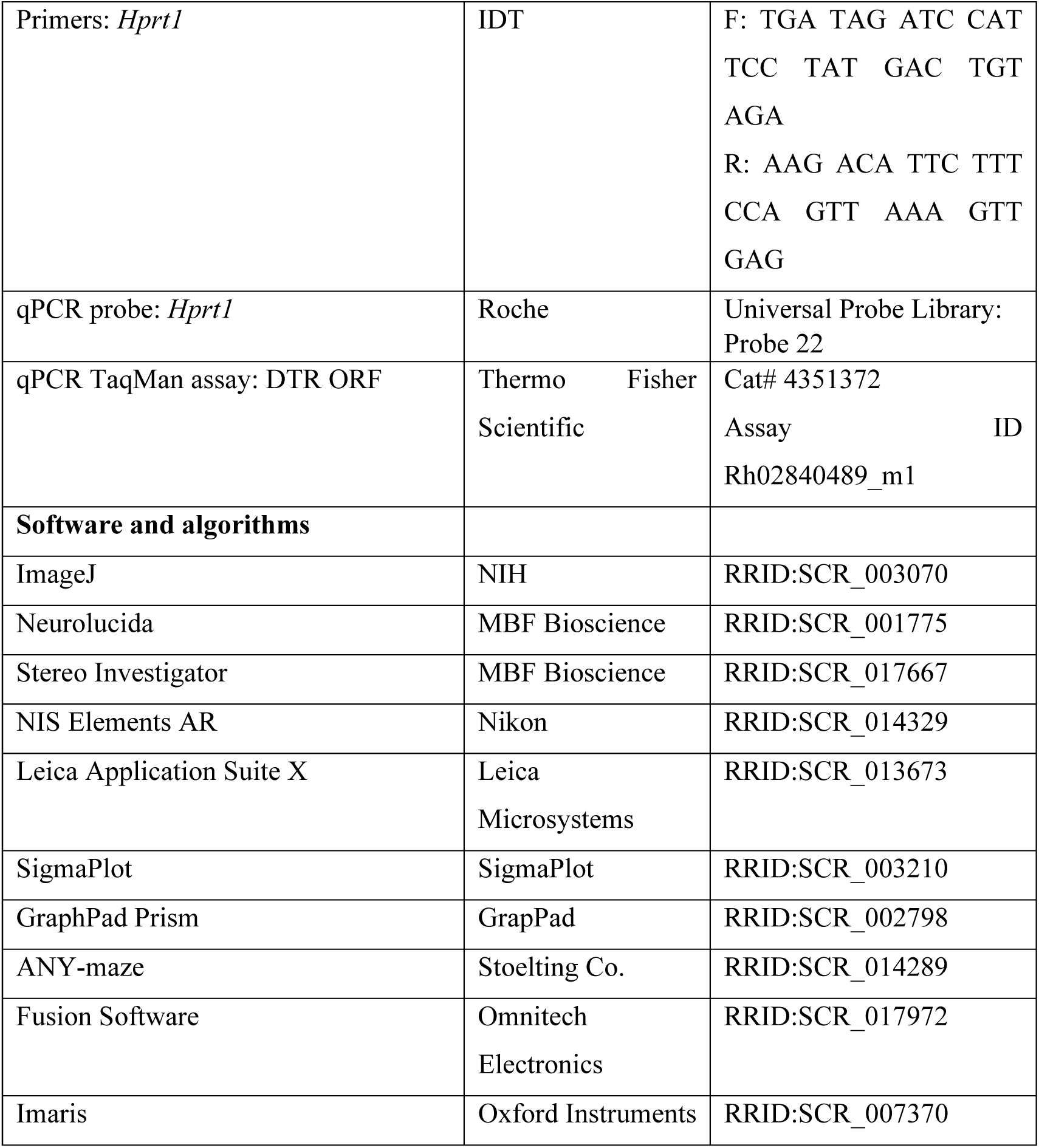
Resources and reagents used in the study.

## Notes

### Competing Interest Statement

The authors have declared no competing interest.

